# Dual function of ERH in primary miRNA biogenesis

**DOI:** 10.1101/2025.09.23.678008

**Authors:** Simon Aschenwald, Aswini Kumar Panda, Theresa Wurzer, Sonja Baumgärtner, Valerie Sophie Ertl, Jakob Hatzer, Andreas Villunger, Sebastian Falk, Sebastian Herzog

## Abstract

MicroRNAs are small non-coding RNAs that mediate post-transcriptional silencing of most mammalian genes. They are generated in a multi-step process initiated by the Microprocessor, a protein complex composed of DROSHA and DGCR8. Recent studies have described the phenomenon of “cluster assistance”, in which a prototypic primary miRNA hairpin can license the Microprocessor-mediated processing of a clustered suboptimal hairpin in *cis*. Genetic screening and mechanistic analyses led to the identification of two critical factors for this process, SAFB2 (scaffold attachment factor B2) and ERH (enhancer of rudimentary homolog), which have been shown to associate with the N-termini of DROSHA and DGCR8, respectively, but also form a complex with each other. However, it remains unclear how SAFB2 and ERH can alter the Microprocessor substrate specificity, and whether the described protein-protein interactions are required for cluster assistance.

In this study, we focused on the role of ERH and show that its loss largely phenocopies the effect of SAFB1/2 deletion on the miRNA transcriptome, suggesting that both factors are involved in the same processes of primary miRNA biogenesis. In this context, our data demonstrate that both SAFB1/2 and ERH are required for efficient Microprocessor feedback regulation via processing of pri-miR-1306, uncovering a clear physiological function of cluster assistance. Mechanistically, our data show that ERH-mediated cluster assistance depends neither on its direct association with SAFB2 nor on its described interaction with DGCR8. In contrast, disrupting the ERH binding site within DGCR8 drives the processing of a subset of cluster assistance-unrelated pri-miRNAs. Thus, this study reveals dual roles of ERH in primary miRNA biogenesis, a largely suppressive one driven by its direct binding to DGCR8, and the other in cluster assistance that does not require DGCR8 binding.

**Highlights:**

- Loss of SAFB1/2 and ERH, respectively, induces overlapping, but not identical defects in primary miRNA biogenesis.
- Both SAFB2 and ERH are involved in Microprocessor feedback regulation.
- ERH-mediated cluster assistance functions independent of its binding to the DGCR8 N-terminus.
- Binding of ERH to the DGCR8 N-terminus confers a largely inhibitory function for primary miRNA biogenesis

## Introduction

MicroRNAs (miRNAs) are small non-coding RNAs of about 21-25 nucleotides (nt) that sequence-specifically repress their target genes and form an additional layer of post-transcriptional gene regulation^1^. In canonical miRNA biogenesis, miRNA genes are transcribed by RNA polymerase II, which generates long primary transcripts (pri-miRNAs) harboring one or several stem-loop structures that encompass the mature miRNAs. These stem-loops are recognized by the nuclear Microprocessor, a protein complex composed of the RNase III DROSHA and two subunits of its co-factor DGCR8 (DiGeorge syndrome critical region 8 homolog; Refs. 2-6). Microprocessor-mediated pri-miRNA cleavage near the base of the stem-loop releases a precursor miRNA (pre-miRNA) that is shuttled into the cytoplasm by RanGTP-dependent Exportin-5, where the RNase III-type endonuclease DICER cleaves off the apical loop to generate an RNA duplex that is incorporated into proteins of the Argonaute (AGO) family^7–10^. The removal of the passenger strand finalizes the formation of the mature RISC (RNA-induced silencing complex), which is the effector complex of miRNA-mediated gene regulation^11,12^. The remaining guide strand directs the RISC to its target sites in mRNAs, which is driven by the miRNA seed sequence as well as by miRNA 3’ complementary base pairing^13–15^. Binding of the RISC to mRNAs, generally to the 3’ untranslated region of transcripts, results in gene repression by translational inhibition and mRNA degradation^16–19^.

In its central role at the initiation of miRNA biogenesis, the Microprocessor complex functions as a gatekeeper whose activity affects all downstream steps and regulatory circuits. As such, the recognition and cleavage of pri-miRNA stem-loops needs to be rigorously controlled to avoid any deleterious consequences for the cell. It has to be ensured that Microprocessor activity is restricted to authentic pri-miRNA stem-loops, i.e. it must not cleave any of the thousands of pri-miRNA-like RNA structures that are predicted to form within the transcriptome simply by chance. Furthermore, it is critical that the processing of the stem-loop itself is very precise, as the cleavage sites define the mature miRNA sequence and, consequently, the target space of the miRNAs. These two prerequisites are met by a combination of distinct pri-miRNA features and sequence motifs that are either necessary for efficient processing or enhance canonical Microprocessor-mediated processing in a modular manner^20–23^. Importantly, many human pri-miRNAs lack some or several of these traits^24^, but are nevertheless recognized and efficiently cleaved. This suggests that additional factors may contribute to proper pri-miRNA processing. Indeed, numerous accessory proteins have been reported to bind directly to the Microprocessor core components or to the pri-miRNA stem-loop structure^2,25–28^, thereby influencing pri-miRNA substrate recognition and/or cleavage efficiency.

While most pri-miRNAs are well defined by the interplay of structural features, sequence motifs and accessory factors, we and others have recently described a novel mechanism that can significantly extend the substrate specificity of the Microprocessor^29–33^. In this process, which we have termed “cluster assistance”, a prototypic pri-miRNA stem-loop (the “helper”) functions as a cis-regulatory element that can license or enhance the processing of neighboring suboptimal pri-miRNAs (the “recipients”) within polycistronic clusters. In consequence, even pri-miRNAs with features that would normally counteract or severely compromise their processing can be recognized and cleaved in a clustered setting. Despite recent advances, we are far from understanding the molecular mechanism of cluster assistance. Current data support a two-stage model in which the Microprocessor is first recruited to the primary miRNA via the helper stem-loop^33^, followed by an ill-defined “transfer” of the DROSHA/DGCR8 complex to the neighboring recipient hairpin in *cis*^31,32^. Of note, an affinity-purified Microprocessor failed to mediate cluster assistance *in vitro* unless supplemented with the total cellular lysate of DGCR8/DROSHA knockout cells^32^, suggesting that additional factors are critical for the processing of the suboptimal stem-loop. Indeed, in our previous work we have identified two factors, SAFB2 (scaffold attachment factor B2) and ERH (enhancer of rudimentary homologue), as critical for cluster assistance^31^, the latter being confirmed by two independent studies^32,34^.

SAFB2, together with the highly similar SAFB1 and the more distantly related SLTM (SAFB-like transcriptional modulator), is part of an evolutionarily conserved family of proteins with DNA- and RNA-binding capability^35^. Along this line, SAFB proteins have been implicated in a number of nucleic acid-centric processes including chromatin organization and genome surveillance, transcriptional regulation as well as RNA splicing and turnover^36–43^. ERH, on the other hand, lacks a canonical RNA binding domain; however, recent data suggests that it likely functions as a scaffold protein that facilitates RNA-binding protein complex formations relevant for multiple mRNA/RNA life cycle steps^44^. Similar to the SAFB family, ERH has been associated with heterochromatin repression and gene silencing, transcription, splicing, RNA processing and nuclear export of mRNAs^44–49^.

At the moment, it remains unclear how these two factors affect the substrate specificity of the Microprocessor, and at which stage of cluster assistance they actually function. Intriguingly, SAFB2 and ERH have been shown to bind to the disordered N-termini of DROSHA and DGCR8, respectively^31,34^, raising the possibility that they may directly enable the Microprocessor to accommodate and cleave suboptimal pri-miRNAs. However, SAFB proteins and ERH have also been demonstrated to interact with each other^50^, but it remains to be determined whether formation of such complexes is relevant in the context of cluster assistance.

Here, we investigate the role of ERH in miRNA biogenesis in more detail, and show that its loss provokes a largely overlapping, but not identical phenotype to SAFB1/2 deficiency, suggesting that both factors are critically involved in the same processes. In this context, our data reveal that both SAFB1/2 and ERH, through cleavage of the DGCR8 mRNA-embedded pri-miR-1306, are required for efficient Microprocessor feedback regulation, uncovering a clear functional role of cluster assistance under physiological conditions. Mechanistically, we show that cleavage of suboptimal cluster assistance recipients depends on both SAFB1/2 and ERH even when the Microprocessor is artificially tethered to its substrate, suggesting their direct involvement in the processing itself. In an attempt to gain mechanistic insight into the role of ERH in miRNA biogenesis, we furthermore tested the functional relevance of its reported direct associations with SAFB2 and DGCR8. With respect to the former, we determined the crystal structure of the ERH-SAFB2 complex, revealing the interaction interface between ERH and SAFB2. Of note, we show that mutation of critical residues compromises complex formation but preserves cluster assistance, suggesting that SAFB2 and ERH fulfill separate roles in this process. Surprisingly, we also find cluster assistance functions independently of ERH binding to DGCR8, and instead show that disruption of the respective interaction motif in DGCR8 enhances expression of a panel of pri-miRNAs that do not depend on cluster assistance. Hence, ERH exerts two distinct roles in primary miRNA biogenesis: One mediated by its direct binding to the Microprocessor, which appears largely suppressive in terms of miRNA biogenesis, and one that is independent of binding to DGCR8 in cluster assistance.

## Materials and Methods

### Plasmid construction

For bacterial protein expression, the genes encoding *Homo sapiens* ERH and SAFB2, together with their truncation variants, were cloned into modified pET vectors. ERH was fused to an N-terminal His_6_–His6-StrepII-Thioredoxin A (TRXA) tag followed by a 3C protease cleavage site. The ERH-binding motif (EBM) of SAFB2 (residues *567-588,* named SAFB2^EBM^) was fused to an N-terminal His_10_–maltose-binding protein (MBP) tag with a 3C site. For the expression of *Homo sapiens* SAFB2, ERH and DGCR8 variants in mammalian cells, the respective cDNAs were cloned into the pMIG-derived pMITHy1.2 or pMIhCD2, in which the GFP cDNA was exchanged for the one encoding cell surface markers Thy1.2 or human CD2. Since the nuclear localization signal (NLS) of DGCR8 has been experimentally placed in its N-terminal region^51^, DGCR8 variants with altered N-termini were expressed with an additional NLS. All amino acid (aa) sequences are available in Table S1.

BoxB constructs for the mutated variants of miR-15a and −181b were generated based on the RNA motif plasmid (Addgene #107253, kindly provided by Paul Khavari) of the RaPID system^52^, and were expressed with the BoxB-miRNA cassette placed in the 3’-UTR of a cDNA encoding a intracellular tail-deletion mutant of the cell surface marker CD8. Single-guide (sgRNA) constructs were designed using CRISPOR or based on the VBC score, and expressed from lentiCRISPR v2 (Addgene #52961, kindly provided by Feng Zhang)^53–55^. Small hairpin RNA (shRNA) vectors for the knockdown of ERH and the respective non-targeting controls were constructed on the basis of human miR-30a, and cloned into an optimized version of the LMP backbone^56,57^. Prime editing guide RNAs (pegRNAs) used the “flip and extension” scaffold modification previously shown to enhance Cas9 activity^58^, and were expressed from the pU-pegRNA GG acceptor plasmid^59^ (Addgene #132777, kindly provided by David Liu) with a tevopreQ1 3’ structural motif for enhanced stability (epegRNA)^60^. All relevant DNA and RNA sequences are available in Table S1.

### Bacterial protein expression and purification

Proteins were expressed in *Escherichia coli* BL21(DE3) derivative strains grown in Terrific Broth at 37°C. When cultures reached an optical density of 2–3 at 600 nm, the temperature was reduced to 18°C. After 2 h of temperature adaptation, protein expression was induced with 0.2 mM isopropyl-β-D-1-thiogalactopyranoside (IPTG) for 12–16 h at 18°C. To purify ERH and SAFB2, cell pellets were resuspended in lysis buffer (25 mM Tris-HCl, 50 mM Na_2_HPO_4_, 500 mM NaCl, 10% (v/v) glycerol, 5 mM β-mercaptoethanol (pH 7.5)) and lysed by sonication. Clarified lysates were applied to a Ni^2+^-charged HisTrap FF column (Cytiva), which was equilibrated in the same buffer. The column was washed with 20 column volumes of lysis buffer, and bound proteins were eluted with 20 mM Tris-HCl, 150 mM NaCl, 500 mM imidazole, 10% (v/v) glycerol, and 5 mM β-mercaptoethanol (pH 7.5).

His-fusion tags were removed by overnight dialysis against 20 mM Tris-HCl, 150 mM NaCl, 10% (v/v) glycerol, and 5 mM β-mercaptoethanol (pH 7.5) at 4°C in the presence of His-tagged 3C protease. A second immobilized-metal affinity chromatography step removed cleaved tags and His-tagged 3C protease. The final purification was performed by size-exclusion chromatography on a Superdex 75 Increase 10/300 column (GE Healthcare) equilibrated in 20 mM Tris-HCl, 150 mM NaCl, and 2 mM dithiothreitol (pH 7.5). All procedures were carried out on ice or at 4 °C.

### Crystallisation, data collection and processing

Crystallization experiments were performed using 5.7 mg/ml ERH with a 1.4-fold molar excess of SAFB2^EBM^ in SEC buffer, employing a vapor diffusion setup. The sample and crystallization solution were mixed in ratios of 1:1 and 2:1 at 22°C. The crystals that yielded the diffraction data for structure determination grew in well B4 of the Morpheus screen (Molecular Dimensions, US), which contained 0.1 M Imidazole, MES monohydrate pH 6.5, 0.03 M Sodium fluoride, 0.03 M Sodium bromide, 0.03 M Sodium iodide, 12.5% (v/v) MPD, 12.5% (v/v) PEG 1000, and 12.5% (w/v) PEG 3350. Crystals from this condition did not need additional cryoprotection and were directly frozen in liquid nitrogen before data collection at 100 K.

Diffraction images were collected at the European Synchrotron Radiation Facility (ESRF, Grenoble, France) at beamline ID23-2 (https://doi.esrf.fr/10.15151/ESRF-ES-771318652). Indexing, integration, and scaling were performed using the xia2 DIALS pipeline^61,62^. Phases were obtained by molecular replacement in Phaser^63^, executed within Phenix^64^, using the Homo sapiens ERH structure (PDB 2NML) as the search model. Automated model building was performed with ModelCraft^65^ via the CCP4 Cloud interface^66^. The model was iteratively completed in COOT^67^ and refined with phenix.refine^68^ and REFMAC5^69^. Model quality was assessed with MolProbity^70^ and PDB-REDO^71^. Data collection and refinement statistics are provided in Table S2.

### Cell lines and cell manipulation

HEK293T (DSMZ #ACC635) and Baf3 (DSMZ #ACC300) cells were cultured in IMDM or RPMI medium, respectively, containing 7.5% FCS, 100 U/ml penicillin and 100 U/ml streptomycin (all Sigma). For Baf3 cells, the medium was supplemented with IL-3-containing supernatant derived from WEHI-3 cells^72^. SAFB1/2-deficient RAMOS cells and their respective controls have been described elsewhere^31^.

For the production of retro- or lentiviral particles, the respective plasmids were transiently transfected into HEK293T cells using branched PEI (polyethylenimine, Polysciences) in a 3:1 (PEI:DNA) ratio, together with packaging vectors pVSVg and either HIT60 or pSPAX2, respectively. Virus supernatants were harvested after 36 to 48 hrs, mixed with Synperonic F108 (Sigma, final conc. 1 mg/ml) and used in spin infections (400 x g) of target cells at 32°C for 90 min.

To generate clones, cells were either plated in limited dilution on 96-well plates or in normal growth medium containing 1% methylcellulose (Sigma; 4000 cP), and colonies were picked and expanded once visible by eye.

### Prime editing

For prime editing, the two epegRNA constructs together with plasmids encoding PE7^73^ (Addgene #214812; kindly provided by Brittany Adamson) and dsRed as a marker were transiently transfected into HEK293T cells, followed by sorting for the highest expression of the components based on dsRed fluorescence after 48 hrs. The sorted cells were singularized, and clones with the edited genomic sequence on both alleles as well as control clones that lacked the edit - but had been subjected to the same treatment - were selected and subsequently sequenced based on a genomic DNA PCR with primers spanning the deletion.

### Small RNA sequencing

Libraries for Illumina sequencing were prepared with the QIAGEN QIAseq miRNA Library Kit (QIAGEN), using 100 ng total RNA isolated with TRIZOL reagent (ThermoFisher) and spiked with reference RNAs (Table S1) as input. The samples were individually indexed using barcoded reverse primers, pooled and sequenced on Novaseq6000 instruments in paired-end mode at the Biomedical Sequencing Facility (BSF) of the Research Center for Molecular Medicine (CeMM), Vienna. Base calls provided by the Illumina Real-Time Analysis (RTA) software were subsequently converted into unaligned BAM format for long-term archival and de-multiplexed into sample-specific BAM files via custom programs based on Picard tools (http://broadinstitute.github.io/picard/). For data analysis, BAM files were converted into FASTQ files using Samtools, and adaptor trimming, mapping against a reference of all mature miRNAs and read counting were done using miRDeep2^74,75^ on the Galaxy server^76^. Spike-in reference RNA counts were retrieved using BBMap^77^. From the resulting matrix, read counts were normalized based on the reference spike-in RNAs and differential expression analysis of the respective samples was then performed using Limma-Voom^78,79^.

### Quantitative RT-PCR

For quantification of mRNAs, total RNA isolated with TRIZOL or the Monarch Total RNA Miniprep Kit (New England Biolabs) was reverse transcribed into cDNA using random hexamers and analyzed by qPCR using either commercial probe-based assays (IDT) or self-designed forward and reverse primers. For precursor and mature miRNA quantification, total RNA was reverse transcribed by self-designed stem-loop assays that use an individual stem-loop primer for cDNA synthesis^80^. The qPCR was then conducted with a specific forward and a universal reverse primer together with a universal probe. All oligonucleotide sequences are available in Table S1. PCRs were run with LUNA qPCR Master Mixes (New England Biolabs). Fold changes were calculated with the 2(−ΔΔC(t)) method, using expression of beta-actin (mRNA) or SNORD68 (miRNA) for normalization.

### Immunoprecipitation, proximity labeling and Western Blotting

For proximity biotinylation, HEK293T cells transiently transfected with plasmids encoding DGCR8, DGCR8-TurboID and ΔN-DGCR8-TurboID (aa261-773; Table S1) were incubated with biotin (Sigma; final conc. 100 µM in normal growth medium) for 1h at 37°C. To stop the labeling reaction, cells were washed four times with ice-cold PBS, detached by pipetting and spun down. Pelleted cells were lysed in ice-cold RIPA buffer (50 mM Tris HCl, pH 7.4, 1% NP-40, 0.25% sodium deoxycholate, 150 mM NaCl, 1 mM EDTA (pH 8)) supplemented with a protease inhibitor cocktail (Roche), and cleared lysates were incubated with 20 µl magnetic streptavidin beads (New England Biolabs; washed twice with RIPA buffer) for 4 h at 4°C. Protein-bound beads were washed once with RIPA buffer, twice with 2% SDS in PBS at room temperature and twice with PBS 0,05% Tween (with incubation of the samples on a turning wheel for 5 min per wash step) before boiling for 6 min in 2x protein loading buffer (125 mM Tris-HCl (pH 6.8), 4% SDS, 20% glycerol, 100 mM DTT, 0.02% w/v bromophenol blue) containing 2 mM biotin.

To analyze the protein interaction between ERH and SAFB2^EBM^, 10 µM and 15µl of purified bait and prey, respectively, were pre-incubated in 200 µL binding buffer (20 mM TRIS/HCl (pH 7.5), 250 mM NaCl, 10% (v/v) glycerol, 0.05% (v/v) IGEPAL, 2 mM DTT) for 30 minutes at 4°C. For MBP pulldowns, samples were added to 35 µL of Amylose resin (New England Biolabs) and for StrepII pulldowns, samples were added to 35 µL of Strep-Tactin XT 4Flow resin (IBA). After the addition of the resin, samples were gently shaken for 1-2 hours at 4°C. Beads were then washed three times with 200 µL binding buffer. Bound material was eluted by incubating the beads for 10 minutes at 4°C with 50 µL binding buffer supplemented with either 10 mM maltose or 50 mM biotin, followed by boiling in protein loading buffer.

For immunoprecipitations out of cellular lysates, HEK293T cells were transiently transfected with the respective constructs and detached by pipetting and spun down after 48 h. Pelleted cells were lysed in ice-cold IP buffer (20 mM Tris-HCl (pH 7.5), 1% Triton X-100, 150 mM NaCl (Figs. 1G and S6B) or 400 mM NaCl (Fig. 1H), 2 mM EDTA, protease inhibitors) and sonicated. Cleared lysates were incubated with 15 µl magnetic anti-FLAG beads (Sigma; washed three times with IP buffer) on a turning wheel at 4°C over night, washed four times with the respective IP buffer and boiled for 10 min in 2x protein loading buffer.

**Figure 1:**
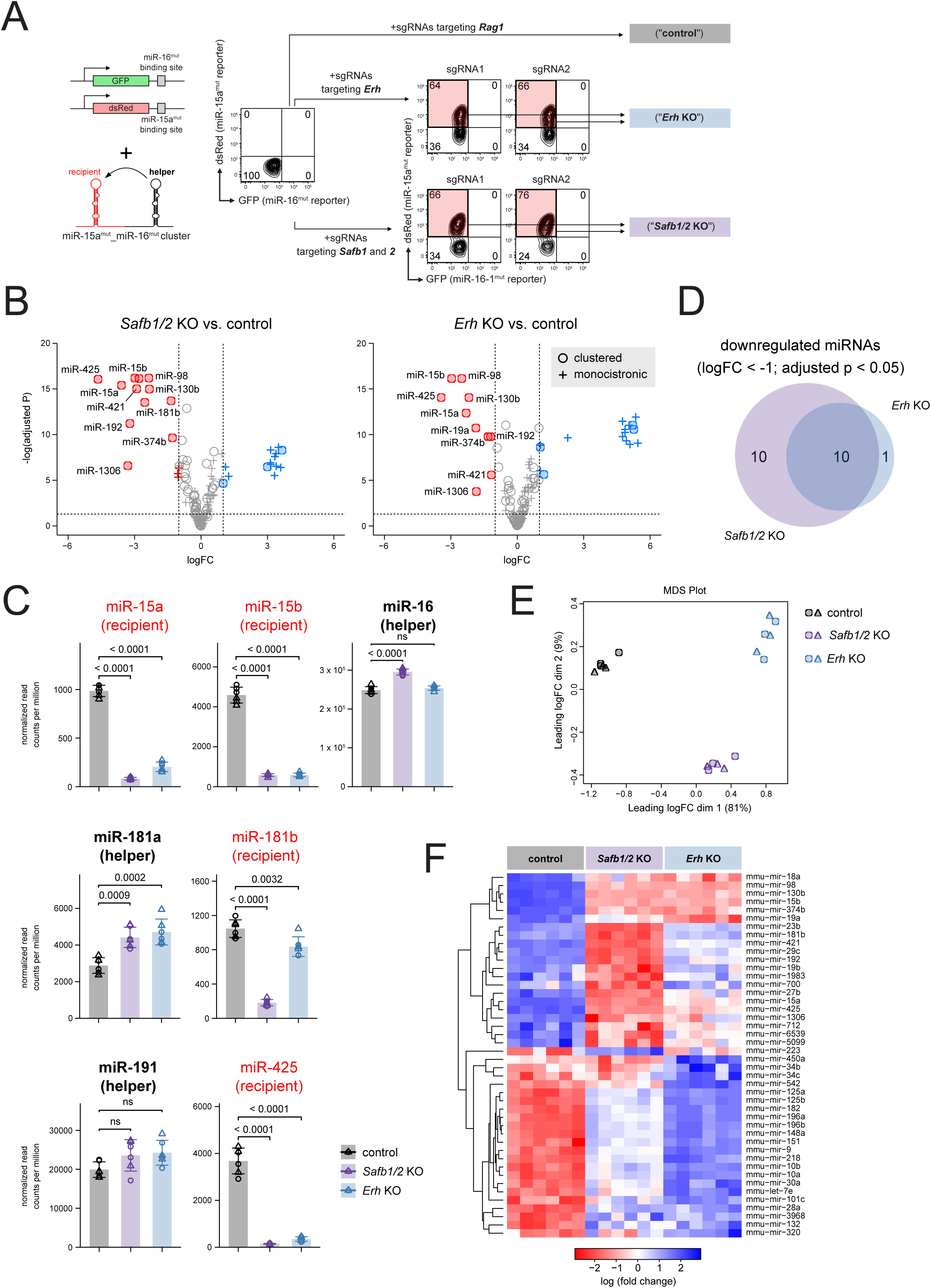
Knockout of *Safb1/2* and *Erh* induces overlapping, but not identical changes in the mature miRNA transcriptome. **A.** Schematic illustration of the dual fluorescence miRNA reporter system expressed in Baf3 screen cells. Parallel expression of the reporters and the mutated cluster results in repression of both dsRed and GFP fluorescence. Subsequent transduction and selection for sgRNAs targeting *Safb1/2* and *Erh*, but not of the unrelated control gene *Rag1*, results in compromised cluster assistance as illustrated by loss of dsRed repression. Colored subsets can be sorted and give rise to bulk/polyclonal KO populations. **B**. Volcano plots comparing mature miRNA expression of bulk *Safb1/2* and *Erh* KO to control cells. Vertical dashed lines mark a log-fold change of “-1” and “1”, respectively, the horizontal line an adjusted p-value <0.05. MiRNAs down- and upregulated in the KO cells are depicted with red or blue circles (for clustered miRNAs with a neighbor on the same strand in less than 2 kb distance) or crosses (for monocistronic miRNAs), respectively. Except for miR-1306, only miRNAs with an average log_2_ CPM (counts per million) > 5 across all samples are plotted. **C**. Bar charts depicting the normalized read counts of mature miRNA species expressed from the miR-15a/b-miR-16, the miR-181a-miR-181b and the miR-191-miR-425 clusters. Helper hairpins are labeled in black, recipients in red. Columns depict mean read counts, error bars display the SD. Data points are visualized as colored circles and triangles for the individual sgRNAs used (n=3+3), numbers indicate p-values. **D**. Venn diagram displaying the overlap of downregulated genes (logFC < −1; adjusted p < 0.05) in the depicted genotypes compared to the control. **E**. 2-dimensional dimensional MDS plot as calculated by limma voom, with individual samples (n=6 for each genotype) depicted as colored circles and triangles depending on the sgRNA used. **F**. Heat map of a selected set of differentially expressed miRNAs in *Safb1/2*, *Erh* and control cells, clustered according to similarity.

Samples prepared as above were subjected to SDS-PAGE followed by Coomassie staining and/or western blotting onto nitrocellulose or PVDF membranes. Primary antibodies for protein detection are listed in Table S1. Immunoreactive proteins were visualized with HRP-labeled secondary antibodies and the ECL system (Advansta) on light-sensitive film or using fluorescent secondary antibodies on the Odyssey CLx (LICOR).

### Flow cytometry

Single-cell suspensions were stained using anti-CD8a, anti-Thy1.2 and, if required, anti-hCD2, followed by streptavidin-BV421. All antibodies and related reagents are listed in Table S1. Data were acquired on an LSR Fortessa or an Aria Sorter (Becton Dickinson).

### Software

Bar charts and volcano plots were generated with RStudio (Posit) using the tidyplots package^81^. Flow cytometric data were analyzed and visualized using Flowjo (Becton Dickinson). Figures were prepared with Affinity Designer (Serif). RNA secondary structures were predicted with the RNAfold web server^82^ and visualized with Varna^83^. All molecular graphics were generated with ChimeraX^84^. Statistical analyses were conducted using Prism 10 (Graphpad).

### Statistical Analysis

Statistical significances was calculated with paired (Figs. 5D and S5C) or unpaired (Figs. 2C, S2B, S2C, 6C, 6D and S6C) t tests when only two groups were compared, or with either one-way ANOVA followed by Tukey post hoc tests (Figs. 1C, S1E, 2B, 2D, 2E, 3D, S3D, 6G and 6H) or one-way repeated measures ANOVA following Bonferroni correction (Figs. 2F, 3B, 3C, S3C, 5C and S5B) for multiple comparisons. Quantitative PCR data are presented as fold changes compared to a control sample to simplify visual interpretation. However, statistical significance was calculated on normally distributed ΔCt values of the respective samples. Error bars represent standard deviation from at least 3 biological replicates.

**Figure 2:**
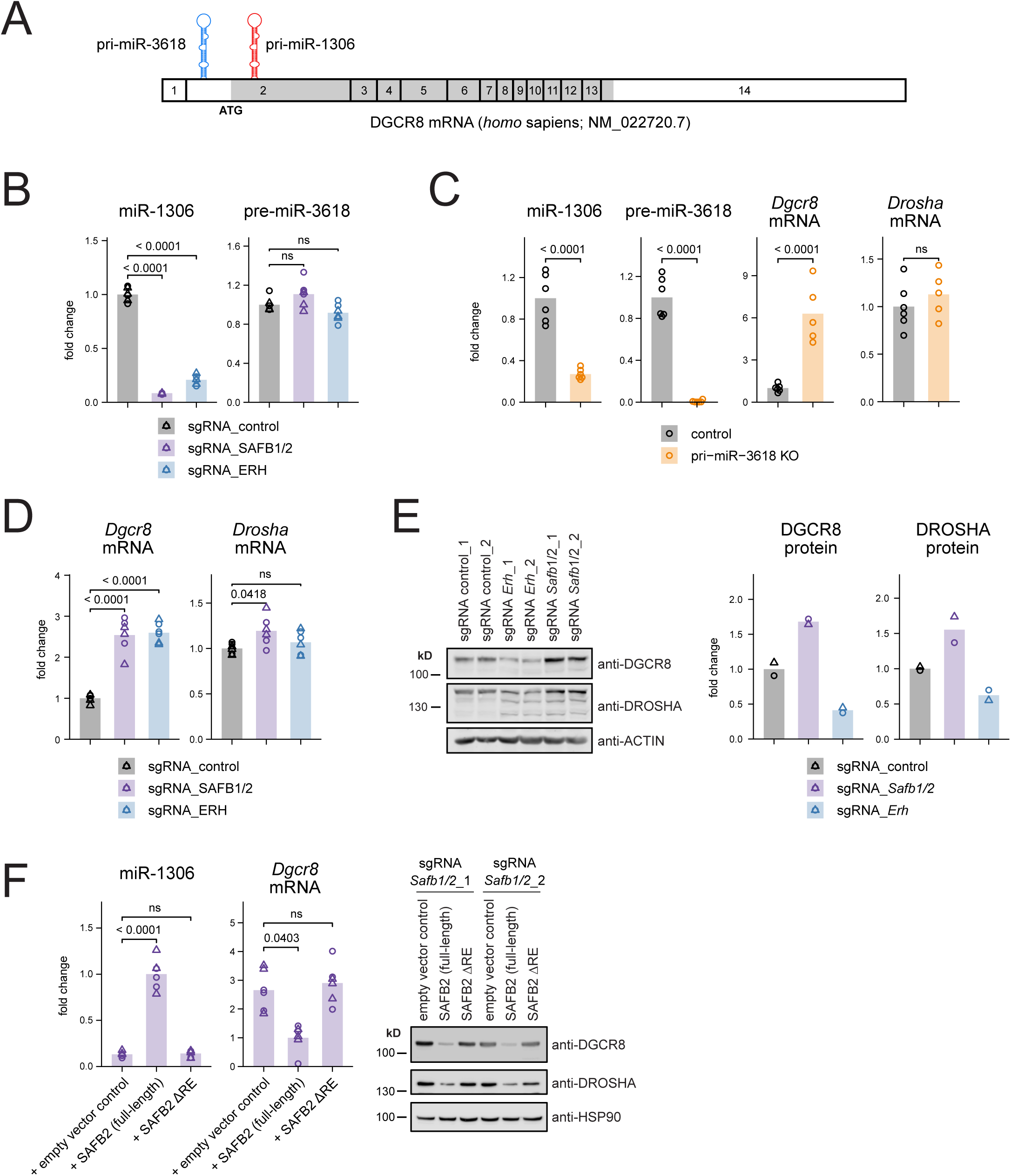
SAFB1/2 and ERH are involved in Microprocessor feedback regulation **A**. Scaled scheme of the human DGCR8 mRNA, with individual exons depicted as numbered boxes. The ORF is marked in dark grey. **B**. Expression of pre-miR-3618 and miR-1306 as quantified by RT-PCR. Individual datapoints are illustrated as colored circles and triangles depending on the sgRNA (n=3+3). **C**. RT-PCR-quantified expression of depicted miRNAs and mRNAs in control and miR-3618 KO cells. **D**, **E**. Expression of DGCR8 and DROSHA mRNAs in SAFB1/2 and ERH KO cells transduced with different sgRNAs, and the corresponding western blot analysis for protein expression and a quantification thereof. **F**. Expression of miR-1306 and DGCR8 RNA as well as protein levels in SAFB1/2 KO cells reconstituted with the depicted SAFB2 variants. Numbers indicate p-values.

**Figure 3:**
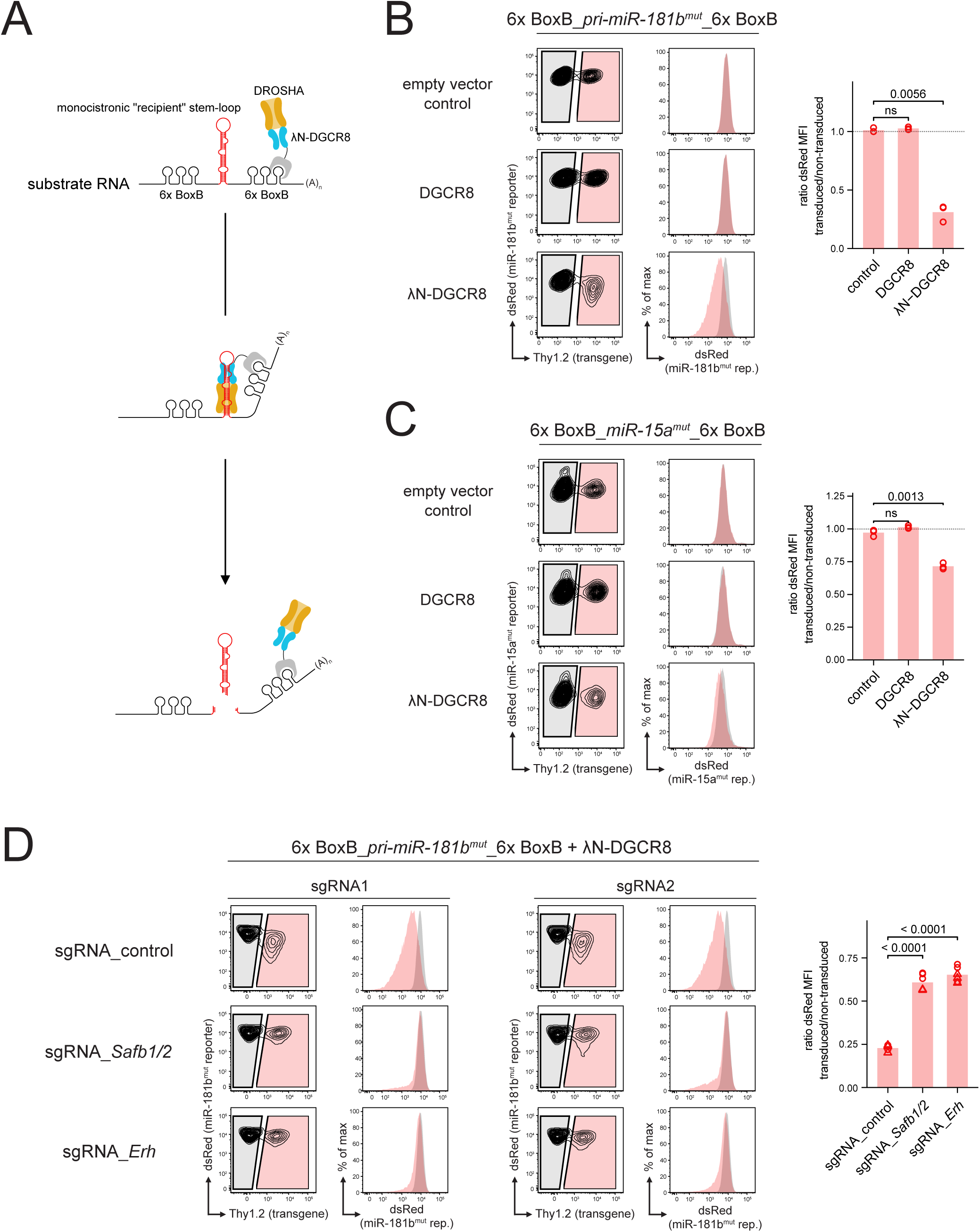
SAFB2 and ERH are directly implicated in the processing of suboptimal pri-miRNA substrates **A**. Schematic illustration of the tethering assay, in which the Microprocessor is recruited to suboptimal substrates via BoxB aptamer-λN interactions. **B**, **C**. Flow cytometric analysis of Baf3 cells expressing the depicted BoxB-miRNA substrate transcripts, the corresponding miRNA reporter and different variants of DGCR8, with Thy1.2 cell surface expression as a marker of the latter. Histograms show overlays of the colored subpopulations. Bar graphs reflecting three independent experiments illustrate the ratio of dsRed-based miRNA reporter expression comparing transduced versus non-transduced cells. **D**. λN-DGCR8 cells as in B demonstrating miRNA reporter repression were transduced with sgRNAs targeting *Safb1/2*, *Erh* and an unrelated control gene. Histogram overlays and the bar graph depict changes in dsRed fluorescence reflecting miRNA expression. Data points are shown as circles and triangles for the individual sgRNAs (n=3+3). Numbers indicate p-values.

## Data availability

Coordinates and structure factors of the ERH/SAFB2^EBM^ complex structure have been deposited in the Protein Data Bank (PDB) with accession code PDB ID 9RMV. Small RNA sequencing data have been deposited under GEO accession numbers GSE311272 and GSE311273, respectively.

## Acknowledgement

We acknowledge the European Synchrotron Radiation Facility (ESRF) for the provision of synchrotron radiation facilities under proposal number MX-2354, and we would like to thank Lindsay McGregor for assistance and support in using beamline ID23-2. This study was supported by the Austrian Science Fund (FWF; grants P30194 and PAT1535423) to S.H..

## Author Contributions

Conceptualization, S.H., S.A., S.F.; Methodology, S.H., S.A., T.W., S.B., V.S.E., J.H.; Formal Analysis, S.H., S.A., A.K.P., T.W., S.B., V.S.E., J.H.; Investigation, S.H., S.A., A.K.P., T.W., S.B., V.S.E., J.H.; Writing – Original Draft, S.H., S.A.; Writing – Review & Editing, S.H., S.A., A.K.P., V.S.E., A.V., S.F.; Visualization, S.H., S.A., A.K.P., S.F.; Supervision, S.H., S.F.; Funding Acquisition, S.H., A.V., S.F..

## Declaration of Interests

The authors declare no competing interests.

## Results

### Small RNAseq analyses of ERH- and SAFB-deficient cells reveal overlapping sets of cluster-assisted miRNAs

While SAFB2 and its closely related family member SAFB1 as well as ERH have been implicated in cluster assistance, their individual contributions have not yet been compared side by side in the same experimental system. To get a general idea about their role in miRNA biogenesis, we generated SAFB1/SAFB2 double-deficient and ERH-deficient cells to assess their miRNA transcriptome by small RNA sequencing. For the generation of the gene knockouts (KOs), murine Baf3 cells emerged as particularly useful, as ERH is largely considered essential^85–87^ and correspondingly its prolonged loss is not tolerated in most cellular systems. In Baf3 cells, however, ERH-deficient cells can be identified, sorted and cultured for extended time, allowing a direct comparison of SAFB1/2 and ERH loss on miRNA expression.

To select for the respective loss-of-function populations, we made use of Cas9^+^ “screen cells” that were initially generated for the CRISPR screen that led to the identification of SAFB2 and ERH (Fig. 1A). These cells express a seed-mutated miR-15a-miR-16-1 cluster (Fig. S1A; hereafter referred to as miR-15^mut^ and miR-16^mut^, respectively), together with dsRed- and GFP-based fluorescent reporters containing the corresponding miR-15a^mut^ and miR-16^mut^ binding sites. Deletion of SAFB1/2 or ERH in this system results in loss of cluster assistance, manifesting as compromised processing of the recipient miR-15a^mut^ and consequently derepression of the respective dsRed reporter signal: At the same time, canonical miRNA biogenesis and thus miR-16 maturation and reporter repression remain unaffected. Of note, expression of full-length SAFB2 in sorted dsRed^+^ SAFB1/2-deficient cells rescued this phenotype (Fig. S1B), and likewise, ERH reconstitution also fully restored dsRed repression in ERH KO cells (S1C).

Thus far, experimental validation of cluster assistance, as evidenced by the concomitant loss of recipient expression upon deletion of the helper, has only been provided for a handful of pri-miRNA candidates, assessed either at their endogenous loci (miR-15a^31^, miR-15b^31^ and miR-451^33^, the latter not being expressed in Baf3 cells) or ectopically expression constructs (e.g. miR-425^31^, miR-181b^31,32^, miR-92a^31^ as well as miR-374a and b^33^). In line with previous reports in heterologous systems ^31,32,34^, small RNA sequencing of the individual populations and of cells expressing sgRNAs targeting a control gene (Rag1) revealed a clear reduction of most of these already confirmed cluster assistance recipients in both genotypes, albeit to varying extent (Figs. 1B and C, Table S2).

In general, the sequencing data showed a remarkable overlap in downregulated and also upregulated miRNAs in SAFB-deficient and ERH-deficient cells (Fig. 1D and S1D), but there were also subtle differences between the genotypes as reflected by the clear separation of the samples in a 2-dimensional MDS (multidimensional scaling) plot (Fig. 1E). SAFB1/2 deficiency, for example, appeared to affect expression of the whole miR-17-92 cluster, whereas the effect of ERH loss was limited to reduced levels of cluster members miR-18a, miR-19a and miR-20a (Fig. S1E). Along the same line, a general analysis revealed a panel of miRNAs that equally depend on SAFB1/2 and ERH, such as miR-98, miR-15b and miR-374b, as well as miRNAs that are predominatly controlled by SAFB1/2 (e.g. miR-23b and miR-192). Overall, the *Safb1/2* knockout seemed to have a more profound effect on the downregulated, potentially cluster assistance-dependent miRNAs, whereas *Erh* knockout resulted in a stronger upregulation of the predominantly monocistronic miRNAs (Fig. 1F). Together, these data strengthen previous reports that link both SAFB1/2 and ERH to cluster assistance, but potentially also indicate their involvement in other processes that shape the mature miRNA transcriptome.

### SAFB2 and ERH regulate Microprocessor levels by targeting the DGCR8 mRNA

One miRNA that was consistently downregulated in both genotypes and also in SAFB-deficient HEK293T and RAMOS cells^31^ was miR-1306, whose stem-loop is embedded about 120 nt downstream of the start codon within the DGCR8 mRNA (Fig. 2A). Together with miR-3618 (encoded on the same exon, but upstream of the ATG), it has been implicated in feedback regulation of DGCR8 by forming a suboptimal Microprocessor substrate that is cleaved depending on the nuclear levels of the complex^88,89^. This is believed to form a rheostat that ensures adequate miRNA levels in all cellular contexts, e.g. throughout development^90^. A detailed analysis of both miR-1306 and pre-miR-3618 (it has been reported that pre-miR-3618 is not further processed by DICER, and thus does not give rise to a mature miRNA^89^) confirmed that the former is expressed dependent on SAFB1/2 and ERH, while pre-miR-3618 was largely unaffected by their absence (Fig. 2B). This suggested a classical cluster assistance setup, with miR-1306 as the recipient and miR-3618 as the helper hairpin. If so, deletion of miR-3618 should markedly reduce expression of miR-1306 due to the loss of cluster assistance, in analogy to the loss of either SAFB1/2 or ERH. Indeed, genomic deletion of miR-3618 by CRISPR/Cas9 resulted in a roughly three-fold reduction in miR-1306 levels (Figs. S2A and 2C). At the same time, it led to a significant increase in *Dgcr8* mRNA levels, indicating the loss of feedback inhibition (Fig. 2C).

Surprisingly, we also found increased *Dgcr8* mRNA and protein levels in SAFB1/2-deficient cells, a phenotype that was confirmed in SAFB-deficient RAMOS cells (Figs. 2D, 2E, S2B and S2C), pointing towards a role of cluster assistance in the Microprocessor rheostat. Accompanying this, DROSHA protein levels were elevated as well, which is in line with reports of its regulation by DGCR8 protein in a yet unknown manner^88^. Upon ERH deficiency, on the other hand, we observed the same increase in *Dgcr8* mRNA levels as demonstrated for the *Safb1/2* knockout cells, but surprisingly, overall DGCR8 and DROSHA protein levels were reduced in Baf3 cells (Figs. 2D and 2E). This suggests a similar mechanism of *Dgcr8* mRNA regulation by SAFB1/2 and ERH, but implies that the latter may play an additional role in DGCR8 protein translation and/or stability. Moreover, we found that reconstitution of SAFB1/2-deficient cells with full-length SAFB2, but not with the cluster assistance-incapable variant lacking the RE-rich domain (ΔRE) spanning amino acid positions 620 to 788, rescued the phenotype on RNA and protein level (Fig. 2F).

Together, these data indicate that cluster assistance-dependent processing of pri-miR-1306, mediated by SAFB proteins as well as ERH, is involved in Microprocessor feedback regulation. As such, in this particular instance, cluster assistance fulfills a clear regulatory role under physiological conditions.

### SAFB proteins and ERH are directly involved in the processing of suboptimal hairpins

While ERH and SAFB proteins are critically involved in cluster assistance, it remains unclear where and how they participate in the process. Based on the discovery of RNA intermediates that indicate sequential processing of the helper and the recipient hairpins, it has been postulated that the helper is primarily required for the recruitment of the Microprocessor to the transcript (Fig. S3A). This recruitment is then followed by helper stem-loop processing and an ill-defined “transfer” of the Microprocessor to the recipient hairpin, which is cleaved in a second step. To investigate whether SAFB and ERH are involved in the initial recruitment and/or transfer of the Microprocessor, or whether they are directly required for processing of the suboptimal cluster assistance recipient, we made use of a system in which the Microprocessor is tethered to the recipient hairpin via BoxB repeats that are efficiently bound by the bacteriophage λ-derived N peptide (λN) fused to the N-terminus of DGCR8 (Ref. 33; Fig. 3A). As such, this experimental setup bypasses the recruitment step and thus isolates the substrate processing itself. Indeed, when combined with fluorescent miRNA activity reporters, tethering of λN-DGCR8 to the two BoxB-flanked monocistronic recipient miRNAs miR-181b^mut^ and miR-15a^mut^, but neither expression of DGCR8 alone nor of pri-miRNAs lacking the flanking BoxB motifs resulted in significant reporter repression (Figs. 3B, C and S3B, C).

Those cells expressing λN-DGCR8 together with the BoxB-flanked pri-miRNAs were then transduced with Cas9 and sgRNAs targeting *Safb1/2*, *Erh* or an unrelated control gene, and dsRed fluorescence was quantified by flow cytometry. Please note that while gene knockout efficiency cannot be easily assessed in this system, the expression of the corresponding constructs in our Baf3 screen cells revealed loss of cluster assistance in about 60 to 80% of transduced cells (Fig. 1A). Here, deletion of either *Safb1/2* or *Erh* significantly reduced dsRed reporter repression for the two BoxB-flanked monocistronic pri-miRNAs, indicating that both factors are directly involved in the processing of suboptimal hairpins under these conditions (Figs. 3D and S3D).

### Mutations targeting the SAFB2/ERH complex interface do not abrogate cluster assistance

To this point, our data for SAFB1/2 and ERH showed striking similarities in terms of regulated miRNAs (Fig. 1) as well as the mode of action for pri-miR-15a and pri-miR-181b cleavage (Fig. 3). Given that a direct interaction between SAFB1/2 and ERH has been reported in the context of phosphorylation of the SR family of splicing factors/regulators, we were wondering whether cluster assistance also depends on a SAFB2/ERH complex or whether both proteins fulfill independent roles (Fig. 4A; Refs. 31, 50). To investigate this, we first characterized the SAFB2/ERH interface *in silico*. In contrast to the initial description^50^, which placed the ERH interaction motif to residues 641-953 for SAFB2, AlphaFold 3 predicted that residues 567–588 (and the corresponding segment in SAFB1) form a short β-hairpin that docks onto each ERH protomer, creating a 2:2 complex. We named this SAFB2 segment the ERH-binding motif (EBM). To verify this prediction, we determined the crystal structure of ERH bound to SAFB2^EBM^ at a resolution of 1.7 Å (Fig. 4B, Supplementary Table S3). The crystal structure obtained is very similar to the model predicted by AlphaFold 3: SAFB2^EBM^ forms a β-hairpin that binds into a groove of ERH formed by α-helices α1-3 and the loop linking α1 and α2. When SAFB2^EBM^ is present, the loop becomes a β-strand, forming a three-stranded antiparallel β-sheet. The interface is stabilized by both hydrophobic and polar contacts. Hydrophobic side chains V572, I583, and V585 of SAFB2 interact with I50 and Y52 in the ERH β-strand and L95 in the α3-helix (Fig. 4C). Multiple polar interactions secure the complex: SAFB2 E569 forms a salt bridge with ERH K34, while SAFB2 R570 is in proximity to two conserved glutamates of ERH, E30 and E37, forming a salt bridge with E37 of ERH. In the hairpin loop, SAFB2 K578 and E580 interact with ERH D53 and S55, respectively, and SAFB2 S584 forms a hydrogen bond with ERH T51 (Fig. 4D).

**Figure 4:**
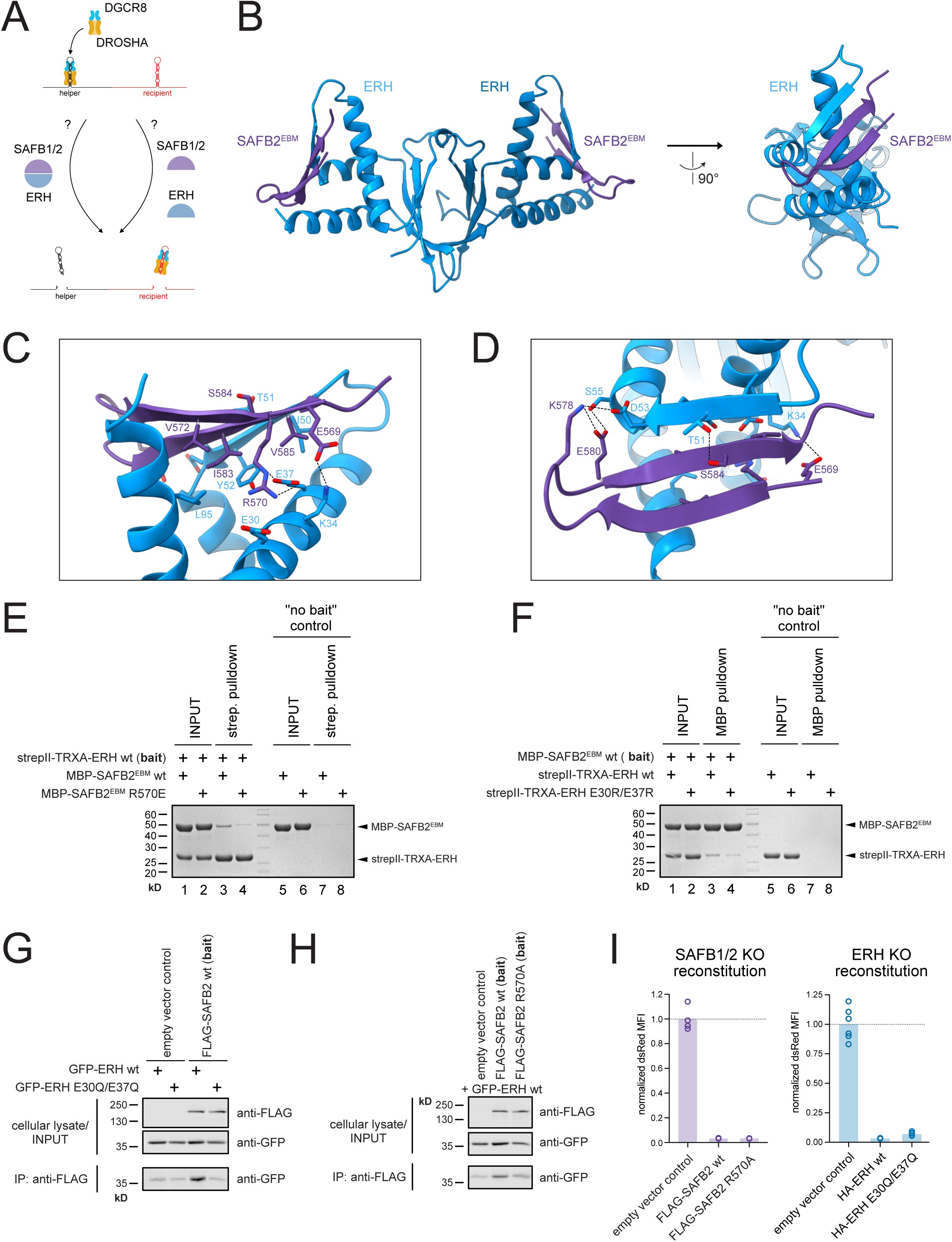
SAFB2 directly associates with ERH through a defined binding motif **A**. Schematic illustration of the models in which cluster assistance is mediated either by a SAFB/ERH complex or by individual functions. **B**. Front and side view of the crystal structure of the 2:2 complex of ERH (blue) with a SAFB2 peptide comprising positions 567–588 (purple). **C**, **D**. Depiction of critical residues involved in SAFB2-ERH interaction. **E**. Coomassie-stained SDS polyacrylamide gels of pulldown experiments with recombinant ERH (bait) and SAFB2^EBM^, respectively. Pulldowns without the bait protein were used as controls. **F**. Reciprocal experiment to E, using SAFB2^EBM^ as the bait. **G**. Pulldown experiments in HEK293T cells transiently transfected with either wt GFP-ERH, GFP-ERH E30Q/E37Q and full-length flag-SAFB2 as a bait. **H**. As G, but using flag-SAFB2 wt and the R570A mutant to pulldown GFP-ERH. **I**. Bar chart of SAFB1/2 (left panel) and ERH knockout screen cells (right panel) reconstituted with the respective constructs. Individual bars depict the mean fluorescent intensity of the dsRed-based miR-15a^mut^ reporter, normalized (dashed line) to the empty vector control. Data points (n=4, n=6) are shown as circles. Numbers indicate p-values.

To confirm the interaction between ERH and SAFB2 observed in the crystal structure, we performed pulldown assays with purified proteins. ERH was produced as a StrepII–thioredoxin A (TRXA) fusion, and the SAFB2^EBM^ peptide was fused to maltose-binding protein (MBP). When StrepII–TRXA–ERH served as bait, it co-precipitated MBP–SAFB2^EBM^; the reciprocal experiment yielded the same result, supporting a direct interaction (Figs. 4E and F, lanes 3). Charge-reversal mutants R570E in SAFB2 or E30R/E37R in ERH weakened the interaction between SAFB2 and ERH, consistent with the interface observed in the crystal structure (Figs. 4E and F, lanes 4). Notably, this validated SAFB2/ERH complex closely resembles the previously reported CIZ1/ERH structure (Fig. S4A to C; Ref. 91).

To confirm the proposed interaction not only with the SAFB2^EBM^ peptide, but also with full-length proteins, we furthermore performed immunoprecipitations in transiently transfected HEK293T cells. Again, wild-type (wt) SAFB2 and ERH pulled down their respective partner, but these interactions were largely abolished with the SAFB2 R570A and ERH E30Q/E37Q variants, respectively (Figs. 4G and H). To evaluate whether the SAFB2/ERH interaction is a prerequisite for cluster assistance, we again made use of the fluorescence reporter system described in Fig. 1A, with miR-15a^mut^ as a prototypic example of a miRNA recipient whose expression depends on a neighboring stem-loop. Deletion of SAFB1/2 or ERH in this system results in loss of cluster assistance, leading to derepression of the respective dsRed reporter signal. As previously shown, expression of full-length SAFB2 in SAFB1/2-deficient and expression of ERH in *Erh* KO cells, respectively, rescued the phenotype (Fig. 4I). Notably, the activity of the interface-disrupting mutants in this assay was comparable to their wt counterparts, indicating that interfering with SAFB2/ERH complex formation does not compromise cluster assistance of pri-miR-15a^mut^ processing.

### ERH mediates cluster assistance independent of its proposed binding site within the DGCR8 N-terminus

Since our findings argued against a critical role of a SAFB2/ERH complex in cluster assistance, we were wondering how else ERH and SAFB1/2 might confer their function. While our own work has placed SAFB2 at the N-terminus of DROSHA^31^, from here on we decided to work on ERH, which has been previously mapped to bind the Microprocessor through an interaction with a motif in the N-terminal region of DGCR8^34^. Confirming this, a DGCR8 mutant lacking this motif (Δ96-125) failed to precipitate GFP-ERH, in contrast to full-length DGCR8 (Fig. 5A). Likewise, proximity biotinylation with DGCR8- and ΔN-DGCR8-TurboID fusions allowed ERH pulldown only in the presence of its proposed DGCR8-binding motif (Fig. S5A), arguing against a clear secondary interaction site.

**Figure 5:**
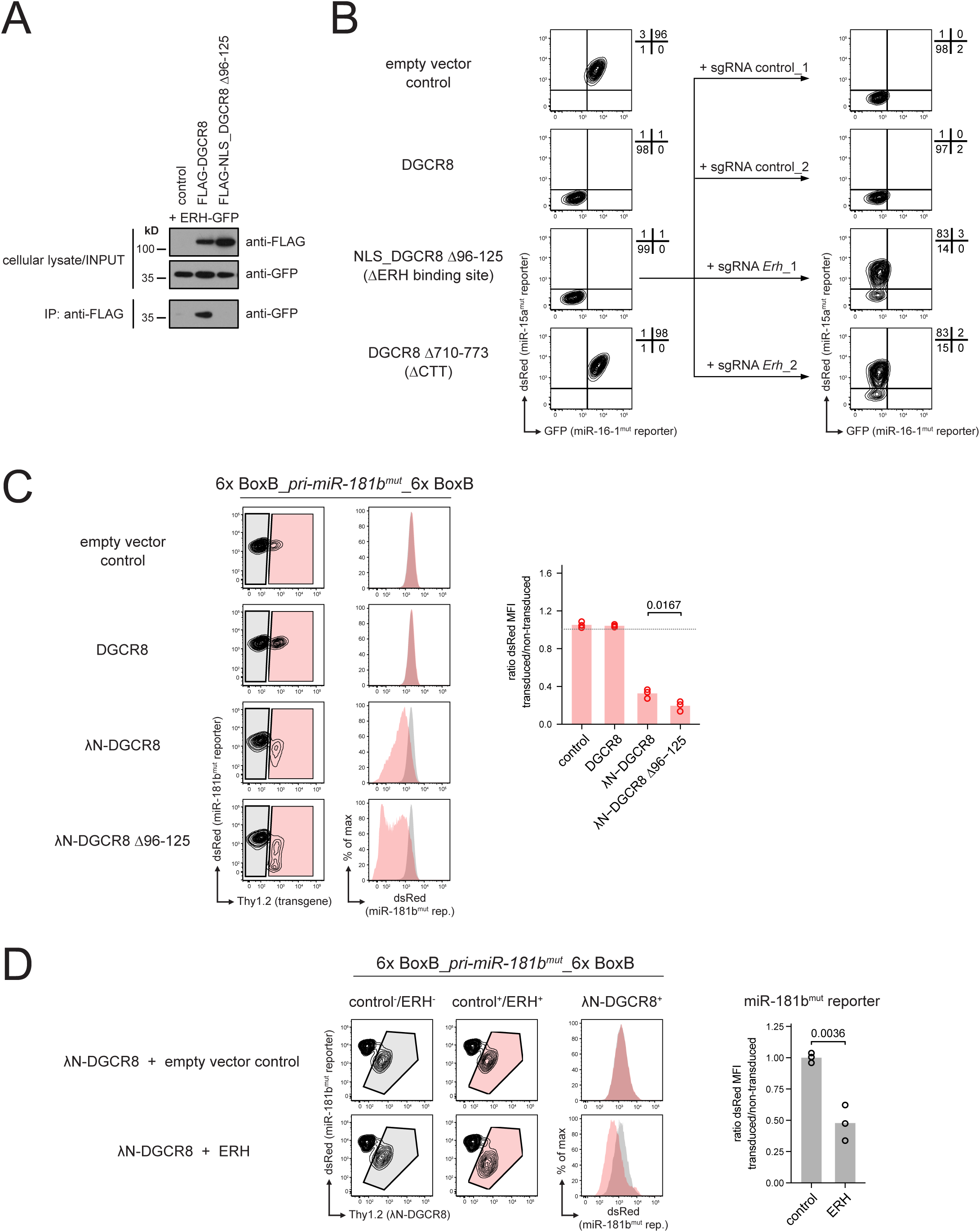
Cluster assistance is independent of ERH binding to its designated DGCR8 motif **A**. Western blot analysis of a pulldown experiment in HEK293T cells expressing GFP-ERH and an ERH binding-competent as well as an incompetent DGCR8 variant (Δ96-125). The GFP-ERH fusion protein is detected with an anti-GFP antibody. **B**. Flow cytometric analysis of DGCR8-deficient dual fluorescent miRNA reporter screen cells reconstituted with different DGCR8 variants as depicted (left panel). DGCR8 Δ96-125-reconstituted cells were further transduced with sgRNAs targeting ERH or a control gene, followed by selection for sgRNA expression and FACS analysis (right panel). Numbers indicate the percentage of cells in the respective quadrants. **C**. Flow cytrometric analysis of Baf3 cells expressing the BoxB-miR-181b^mut^ substrate transcript, the corresponding miRNA reporter and different variants of DGCR8. Histograms show overlays of the colored subpopulations. Bar graphs provide the ratio of dsRed-based miRNA reporter expression comparing transduced versus non-transduced cells for three independent experimental replicates. **D**. Analysis of cells as in C, but with additional expression of a control vector (upper line) or ERH (lower line). Histogram overlay the colored subpopulations, corresponding to λN-DGCR8+ transduced (red) and non-transduced (grey) cells. The bar chart shows the quantification of three independent experiments, comparing the effect of ERH or control vector expression on the dsRed-based miRNA reporter. Numbers indicate p-values.

Accordingly, we anticipated that disruption of the ERH binding site within DGCR8 would phenocopy the loss of cluster assistance observed upon ERH deletion. However, when we reconstituted screen cells deficient for DGCR8, characterized by derepression of both miRNA reporters, with the DGCR8 Δ96-125 variant, canonical miRNA processing (GFP-based miR-16^mut^ reporter) as well as cluster assistance (dsRed-based miR-15a^mut^ reporter) were fully restored (Fig. 5B, left panel). In contrast, control cells and cells expressing a DGCR8 variant lacking the C-terminal tail (Δ710-773), required for DROSHA interaction, failed to do so. Notably, CRISPR/Cas9-mediated knockout of *Erh* in the cells reconstituted with DGCR8 Δ96-125 still induced the derepression of the dsRed-based reporter, suggesting that ERH mediates cluster assistance independent of its described binding site within DGCR8 (Fig. 5B, right panels).

To strengthen this point, we again utilized the Microprocessor recruitment system and tethered λN-DGCR8 Δ96-125 to suboptimal miR-181b^mut^ and miR-15a^mut^, respectively. Based on our reconstitution experiments in DGCR8-deficient cells, we anticipated that recruitment of λN-DGCR8 lacking the ERH binding site would be as potent as its wt variant. Surprisingly, with both reporters, the Δ96-125 mutant appeared to be even more effective in terms of reporter repression (Figs. 5C and S5B). Based on our findings that ERH mediates cluster assistance independent of its binding to DGCR8 (Fig. 5B), we speculated that ERH may have two distinct functions in miRNA biogenesis, one being mediated via its binding to the DGCR8 N-terminus, and one being independent of this interaction. If so, we envisioned that the cellular pool of ERH may be rate-limiting, i.e. that both processes compete for ERH. In this scenario, overexpressed λN-DGCR8 might sequester otherwise free (or at least DGCR8 aa96-125-unbound) ERH and interfere with its use in other cellular pathways. To test this hypothesis, we overexpressed ERH in λN-DGCR8-positive cells and monitored repression of the fluorescent miRNA reporter. Indeed, expression of ERH significantly enhanced reporter repression (Fig. 5D and S5C), analogous to expression of the λN-DGCR8 Δ96-125 mutant. We conclude that ERH can be sequestered by DGCR8 in our tethering assay, thereby limiting its function in cluster assistance and suboptimal pri-miRNA stem-loop processing.

### Dual function of ERH in primary miRNA biogenesis

These findings, of course, raised the question about the functional relevance of ERH binding to the DGCR8 N-terminus. To address this, we used a paired prime-editing approach to disrupt the described ERH binding motif by modifying the endogenous DGCR8 loci in HEK293T cells, thereby avoiding any overexpression artifacts (Ref. 92; Fig. 6A). For technical reasons, we deleted a slightly smaller region of the N-terminus as compared to the previous experiments (Δ96-125 vs. Δ103-126Q), but pull-down as well as tethering experiments demonstrated that this mutant also lacks the capability to interact with ERH (Figs. S6A and S6B). Genomic DNA sequencing, western blot and PCR analysis of selected prime-edited clones as well as the corresponding controls confirmed the expected edits, with DGCR8 running at a slightly lower molecular weight due to deletion of 22 aa (Figs. 6B and C). To our surprise, disruption of the ERH binding site resulted in an increase in DGCR8 protein compared to controls, possibly indicating an impact on Microprocessor feedback regulation. Indeed, we found reduced levels of pre-miR-3618 and miR-1306 in clones lacking the ERH binding site (Fig. 6D), confirming that a DGCR8/ERH complex is required for adequate *DGCR8* mRNA processing and thus Microprocessor expression.

**Figure 6:**
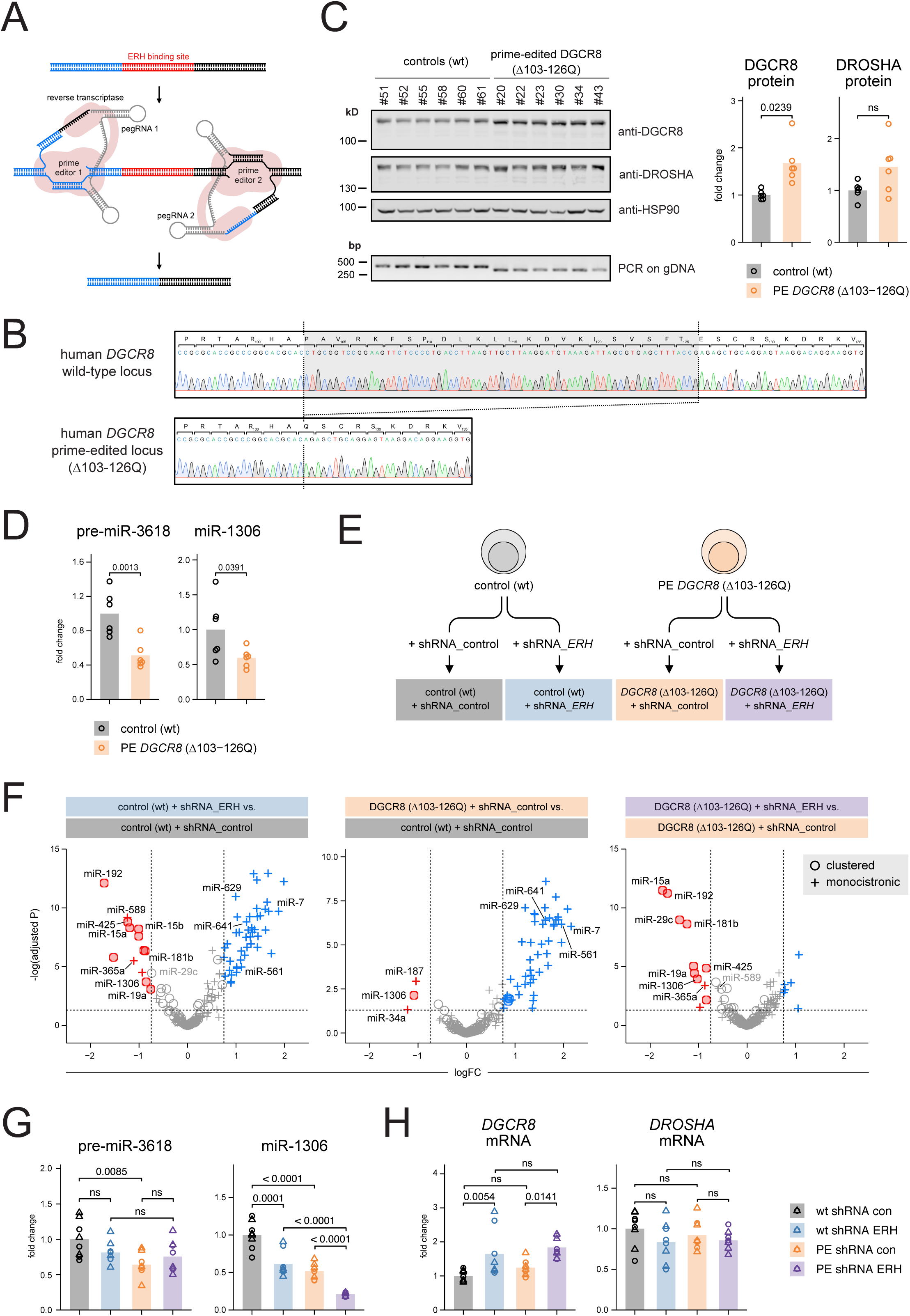
ERH confers two distinct roles in primary miRNA biogenesis **A**. Schematic illustration of the paired prime-editing approach used for precise deletion of the ERH binding site in DGCR8. **B**. Representative Sanger sequencing result of a wt control and a prime-edited DGCR8 Δ103-126Q clone. The corresponding primary amino acid sequence is shown on top. **C**. Western blot and genomic DNA analysis of control and prime-edited HEK293T cells clones. Protein expression of DGCR8 and DROSHA has been plotted on a bar chart as a fold change compared to the control cells, with each clone corresponding to one data point. **D**. Pre-miR-3618 and miR-1306 expression of clones as in C, quantified by RT-PCR. Numbers indicate p-values **E**. Overview of the ERH knockdown experiment. Four clones per genotype were individually transfected with two control shRNAs and two shRNA targeting ERH, resulting in a total of 32 samples. **F**. Volcano plots comparing mature miRNA expression in populations colour-coded as in E. Vertical dashed lines mark a log-fold change of “-0.75” and “0.75”, respectively, the horizontal line an adjusted p-value <0.05. Down- and upregulated miRNAs are depicted with red or blue circles (for clustered specimen, with another miRNA on the same strand in < 2 kb distance) or crosses (for monocistronic miRNAs), respectively. Except for miR-1306, only miRNAs with an average log_2_ CPM > 6 across all samples are plotted. MiRNAs downregulated in the prime-edited cells are depicted as orange circles. **G, H**. Quantitative PCR analysis of depicted precursor and mature miRNAs as well as mRNAs for sample groups as in E. Numbers indicate p-values.

To characterize the phenotype of the ERH binding site mutation in more detail, in particular in comparison to a global loss of ERH expression, we set up an experiment analogous to Fig. 5B. In detail, the prime-edited *DGCR8* Δ103-126Q and the control cells were transiently transfected with two individual shRNAs targeting *ERH*, resulting in strong knockdown of *ERH* mRNA after five days compared to non-targeting control shRNAs (Figs. 6E and S6C). The resulting sample groups were then analyzed by small RNA sequencing and compared side-by-side. In line with previous reports^32,34^, knockdown of ERH on a wt background resulted in reduced expression of confirmed cluster assistance targets miR-15a, miR-15b, miR-181b and miR-425 (Fig. 6F, left panel, Table S4). Beyond those, most of the downregulated miRNAs were again part of miRNA clusters, with the exception of monocistronic miRNAs such as miR-589 and miR-365a, both of which have recently been described to be regulated by ERH in a cluster assistance-independent manner^93^. Notably, we found these two monocistronic miRNAs downregulated also in a dataset of SAFB1/2-deficient RAMOS cells^31^ (Fig. S6D), suggesting that SAFB proteins can promote – in analogy to ERH – primary miRNA biogenesis beyond cluster assistance. Recapitulating our Baf3 *Erh* KO findings, we furthermore identified a panel of predominantly monocistronic miRNAs that became upregulated upon reduced global ERH expression, including miR-7, miR-629, miR-641 and miR-561, pointing towards a negative regulatory role of ERH in the context of distinct miRNAs.

In comparison, disruption of the ERH binding site in DGCR8 Δ103-126Q cells did only result in downregulation of a very small subset of mature miRNAs, but revealed the same pattern of upregulated miRNAs observed upon ERH knockdown (Fig. 6F, center panel). This indicates that the ERH binding site embedded within the DGCR8 N-terminus is dispensable for cluster assistance, and vice versa, that the process is mediated by unbound or by otherwise with DGCR8 and/or DROSHA associated ERH. Supporting this, ERH knockdown in the prime-edited DGCR8 Δ103-126Q cells again reduced expression of prototypic cluster assistance recipients (Fig. 6F, right panel). Of note, a direct comparison of shERH-expressing cells of both genotypes revealed largely identical changes in miRNA biogenesis, with minor differences probably due to incomplete ERH knockdown (Fig. S6E). This implies that the DGCR8 Δ103-126Q modification does not interfere with any miRNA-related processes other than ERH binding.

Together, these data clearly define two distinct functions of ERH in miRNA biogenesis: On the one hand, ERH is critical for cluster assistance through a yet to be identified mechanism that is independent of DGCR8/ERH complex formation. On the other hand, ERH confers a predominantly repressive role that is established by its direct association with the DGCR8 N-terminus. Interestingly, both processes appear to converge on Microprocessor feedback regulation: While most miRNAs affected by DGCR8 Δ103-126Q become upregulated when ERH cannot bind to DGCR8 anymore, miR-1306 is found downregulated in these conditions (Fig. 6F, center panel). Given that cells expressing DGCR8 Δ103-126Q have reduced levels of both miR-1306 and pre-miR-3618 (Fig. 6D), we infer that the loss of the DGCR8/ERH interaction interferes with pri-miR-3618 processing, with the accompanying reduction of miR-1306 being an indirect, secondary effect explained by cluster assistance. Global ERH loss, however, not only targets pri-miR-3618, but also disables cluster assistance and thereby reduces miR-1306 even further. Supporting this hypothesis, only the combination of DGCR8 Δ103-126Q with global knockdown of ERH results in full loss of miR-1306, resulting in even higher DGCR8 mRNA levels than either modification alone (Figs. 6G and H).

## Discussion

Previous experiments demonstrated partially overlapping defects in primary miRNA biogenesis upon SAFB1/2 deletion and ERH knockdown, respectively, suggesting that both either participate in the same pathway, e.g. as a mandatory protein complex, or that they operate in individual processes that are both required for efficient cluster assistance^31–33^. However, the underlying data were derived from different experimental systems and laboratories, making a direct comparison of both components and their individual contribution challenging. Combining a cellular system that tolerates loss of SAFB1/2 and ERH with a fluorescent reporter system for the identification of cells defective for cluster assistance, our small RNA sequencing data presented here reveal the same pattern of downregulated mature miRNAs upon deletion of both factors, albeit with small differences. Of course not all of these effects can be linked to defective cluster assistance, as the differential expression of miRNAs we observe likely reflects direct as well as indirect effects of SAFB1/2 and ERH deletion, and can also be the consequences of events covering all stages of the miRNA lifecycle. However, it is striking that the downregulated miRNAs are predominantly clustered, and it will be interesting to evaluate whether candidates such as miR-130b, miR-98 and miR-192, which have been identified in addition to established recipients such as miR-15a and b, miR-425 and miR-181b, also turn out to depend on cluster assistance.

One SAFB- and ERH-dependent miRNA we found in all our datasets across species and cell lines was miR-1306, which has been implicated in adjusting global Microprocessor levels and downstream miRNA dosage, through Microprocessor feedback regulation^88,89^. Together with pri-miR-3618, pri-miR-1306 is encoded within the second exon of DGCR8, and they fold into two hairpins that flank the start codon on the DGCR8 mRNA. Previous data indicate that pri-miR-3618 is the better Microprocessor substrate, whereas pri-miR-1306 is only poorly processed, which is also reflected by the comparably low read numbers found in small RNA sequencing data (Refs. ^88,89^; our data). In consequence, pri-miR-3618 appears to be the key substrate in feedback regulation, and its deletion results in strong upregulation of *Dgcr8* mRNA and protein levels (Ref. ^90^ and Fig. 2C), a scenario that has even been shown to promote irreversible Microprocessor aggregation. This processing hierarchy can at least in part be explained by our findings that pri-miR-1306 is a recipient of cluster assistance mediated by the helper pri-miR-3618, i.e. pri-miR-1306 is largely cleaved downstream of pri-miR-3618, independently confirming recent findings^93,94^. Accordingly, deletion of SAFB1/2 does not alter the processing of pri-miR-3618, but reduces only the levels of miR-1306. For ERH, the situation is more complex: In the human system, ERH not only mediates recipient processing (pri-miR-1306) via cluster assistance, but at the same time also turns out to be necessary for efficient helper cleavage (pri-miR-3618). In the analyzed murine cells, in contrast, the direct effect of ERH on pri-miR-3618 is rather weak (Fig. 2B), but its loss nevertheless strongly affects miR-1306 and *Dgcr8* mRNA levels. Surprisingly, DGCR8 protein in these cells is reduced despite higher mRNA levels, pointing towards species and/or cell line-specific post-transcriptional consequences of ERH loss.

In order to gain mechanistic insight into the role of SAFB1/2 and ERH in cluster assistance, and considering the overlapping pattern of small RNA expression upon their deletion, we tested whether both factors need to form a complex to confer their function. However, while we have been able to describe the previously reported interaction of SAFB1/2 and ERH in more detail, charge-reversing or -abolishing mutants of critical residues in ERH as well as in SAFB2, which weakened complex stability in pulldown experiments, remained cluster assistance-competent at least in the context of prototypic miR-16-mediated miR-15a processing. It therefore appears unlikely that a stable SAFB2/ERH complex is required for the process, but rather argues for both factors being involved in individual functional modules that converge on and together are critical for cluster assistance. Nevertheless, our structural and biochemical data may provide insight into the general function of ERH, as the SAFB2/ERH interaction describe here closely resembles the previously reported CIZ1/ERH structure^91^. In both cases, a peptide folds into a β-hairpin that docks onto the same surface of ERH, and each hairpin features a prominent arginine that contacts ERH residue E37. Given this similarity, it is tempting to speculate that this mode of interaction, with a β-hairpin docking into the same groove of ERH, is a characteristic feature that is also shared with additional interactors beyond SAFB2 and CIZ1. If so, it is likely that the resulting competition for the ERH binding interface confers additional regulatory roles, particularly in light of ERH being recently described as a protein that forms a dense network of interactions with several RNA-binding proteins^44^.

So how, if not via a complex, do SAFB1/2 and ERH exert these apparently separate yet complementary roles in cluster assistance? Based on their reported association with the N-termini of DROSHA and DGCR8, respectively, we hypothesized that their individual or combined binding induces a conformational shift within the Microprocessor, which may relax structural constraints and enable the cleavage of suboptimal hairpins. However, in line with recent data^94^, our work clearly demonstrates that cluster assistance is not driven by DGCR8-bound ERH, at least not via its described and structurally characterized binding site^95^. Importantly, experiments from the Bartel lab have shown that also an N-terminally truncated DROSHA variant that lacks the described SAFB2 binding site is cluster assistance-competent^32^. This is in accordance with our own data demonstrating a rescue of miR-15a-targeted cluster assistance upon reconstitution of DROSHA-deficient Baf3 screen cells with a DROSHA Δ1-390 mutant (data not shown), and has recently been confirmed^94^. Thus, while SAFB1/2 and ERH can associate with the Microprocessor, their interactions with the described protein domains seem to be dispensable for cluster assistance.

It is certainly possible that one or both of them associate with DROSHA and/or DGCR8 through alternative, yet to be characterized binding motifs. Such a scenario appears unlikely for ERH, as our proximity biotinylation experiments with a DGCR8 mutant lacking the N-terminal region failed to pull down any protein. However, a recent study reported that SAFB2 can be co-immunoprecipitated with DGCR8 even in the absence of DROSHA^94^, suggesting a potentially distinct mode of recruitment that warrants further investigation. Alternatively, we cannot rule out that cluster assistance is mediated without physical interaction of SAFB1/2 and ERH with the Microprocessor. It is tempting to speculate that SAFB proteins and ERH shape or define the nuclear environment in which primary miRNA biogenesis and cluster assistance takes place, in line with their role as adapter proteins that are likely deeply embedded in various protein-protein interaction networks. However, these theories need to be experimentally tested in the future.

Mechanistically, our data indicate that both SAFB1/2 and ERH are required for miRNA activity when DGCR8 is tethered to suboptimal miR-15a and miR-181b, suggesting that both factors are directly involved in recipient processing. Contradicting these findings, however, despite using a similar experimental setup, Jang et al. report that pri-miR-451 processing depends on ERH, but not on SAFB2, when the recruitment step is bypassed^93^. Further complicating the situation, a recent preprint reports the opposite findings for for the same pri-miRNA, albeit with a slightly different tethering approach^94^, finding SAFB proteins to be critical for processing, whereas ERH appears to be dispensable. These discrepancies likely reflect differences in tethering construct design and possibly also in pri-miRNA dependencies, and given the artificial nature of such assays, distinguishing physiological mechanisms from experimental artifacts will be critical.

Nevertheless, by comparing the singular and combined phenotypes of global ERH knockdown and ERH binding site deletion, we have been able to precisely dissect the individual contributions of ERH and demonstrate that its recruitment to DGCR8 clearly impacts Microprocessor function beyond cluster assistance. Except for pri-miR-3818 and two additional candidates, the formation of the DGCR8/ERH complex appears largely inhibitory, as illustrated by upregulation of mature miRNAs such as miR-7, miR-641 and miR-629 upon mutation of the binding motif, respectively. As such, our findings challenge a recently proposed model in which ERH generally enhances Microprocessor activity through binding to DGCR8^93^. In fact, our data define the monocistronic pri-miRNAs most prominently affected by ERH loss in the respective study, pri-miR-589 and pri-miR-365a, as independent of the ERH binding site, but dependent on global ERH and SAFB1/2 (Figs. 6F and S6D). This indicates that cluster assistance and processing of these monocistronic miRNAs follow the same rules, possibly arguing for the same underlying molecular mechanism.

In the context of the ERH/DGCR8 complex, for which structural and biochemical data suggest a 2:2 stoichiometry, we hypothesize that ERH either bridges several Microprocessors or, more likely, brings both DGCR8 N-termini within one Microprocessor complex into close proximity. Despite our limited knowledge of these largely disordered and structurally unresolved regions, it appears feasible that such an induced conformational change in the complex may be transmitted towards DROSHA, thereby altering its substrate binding and/or processing capabilities. Of note, although SAFB2’s critical role in cluster assistance is independent of its binding to the DROSHA N-terminus, it will be interesting to investigate whether a similar regulatory mode also operates on this end of the Microprocessor.

In consequence, we postulate a model in which ERH confers two distinct functions in primary miRNA biogenesis (Fig. S6F): One the one hand, it serves as an auxiliary factor to the Microprocessor through direct association with the binding motif embedded within the DGCR8 N-terminus, which has a predominantly repressive role in Microprocessor function for a subset of miRNA stem-loops. On the other hand, ERH mediates cluster assistance independent of the described DGCR8/ERH complex, in a process that obviously co-depends on SAFB proteins. At the moment, we can only speculate how these two factors can alter the substrate specificity of the Microprocessor. Although the mechanisms by which SAFB1/2 and ERH modulate Microprocessor substrate specificity remain unresolved, our findings provide a framework for future investigations into how these factors enable the phenomenon of cluster assistance.

---

**Figure S1.**
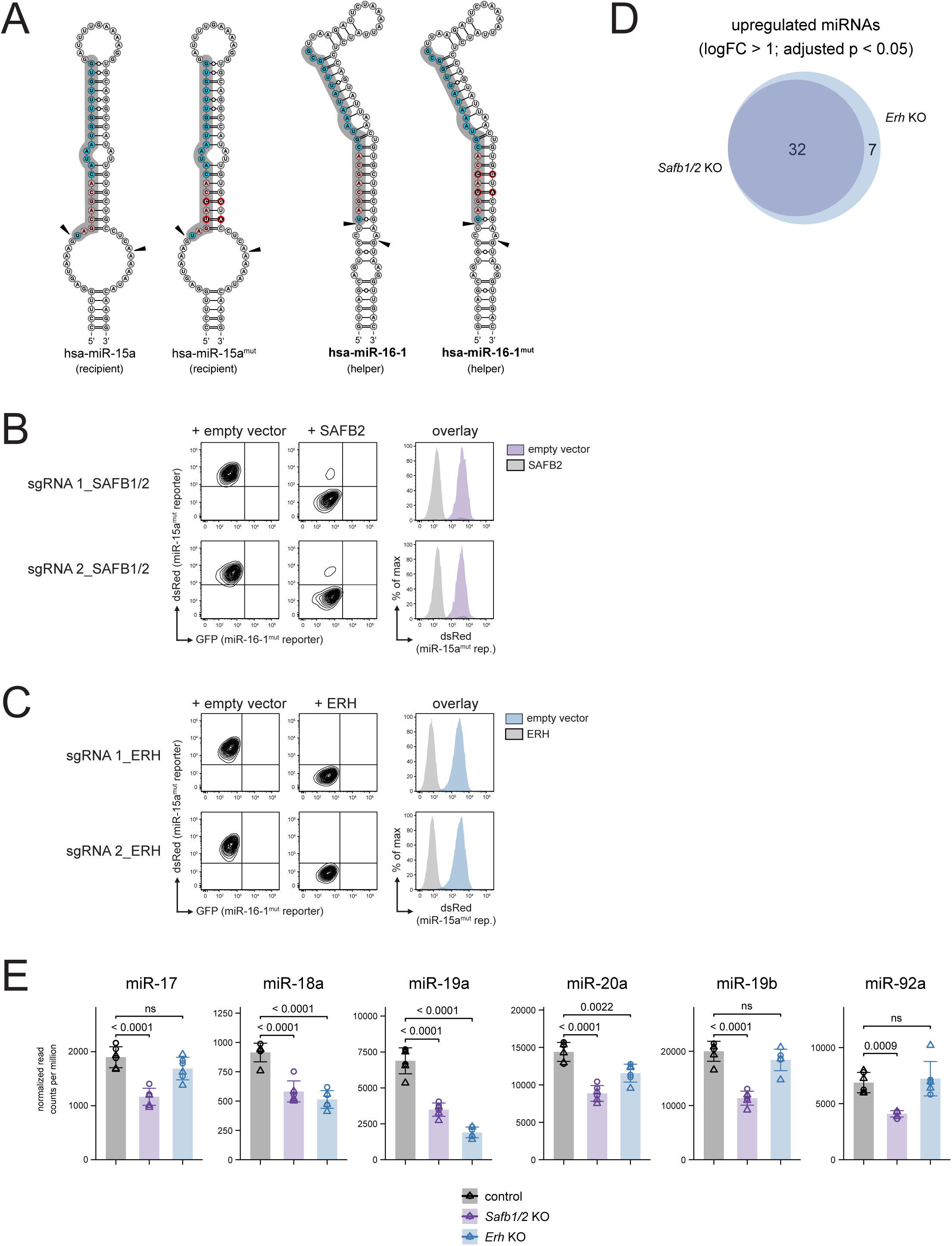
**A**. Predicted structures of miR-15a and miR-16-1 and their respective seed mutants used throughout this study. The mature miRNA sequences within the pri-miRNAs are shown as turquise bases on a gray background, with seed sequences in red and black (mutated). Triangles mark the predominant Microprocessor cleavage sites. **B**, **C**. Representative flow cytometric analysis of *Safb1/2* (B) and *Erh* (C) KO loss-of-function cells expressing the dual fluorescence reporter as in Fig. 1A, characterized by a prominent upregulation of the dsRed reporter due to defective cluster assistance. KO cells were generated with two independent sgRNAs, sorted for dsRed expression and retrovirally transduced with an empty control vector or with human SAFB2 and ERH, respectively. Contour plots show the populations positive for transgenic expression, and histogram overlays illustrate the rescue of reporter repression upon SAFB2 and ERH reconstitution. **D**. Venn diagram displaying the overlap of upregulated genes (logFC < −1; adjusted p < 0.05) in the depicted genotypes compared to the control. **E**. Bar charts depicting the normalized read counts of mature miRNA species expressed from the miR-17-92 cluster in the respective genotypes. Columns depict mean read counts, error bars display the SD. Data points are visualized as colored circles and triangles for the individual sgRNAs used (n=3+3), numbers indicate p-values.

**Figure S2.**
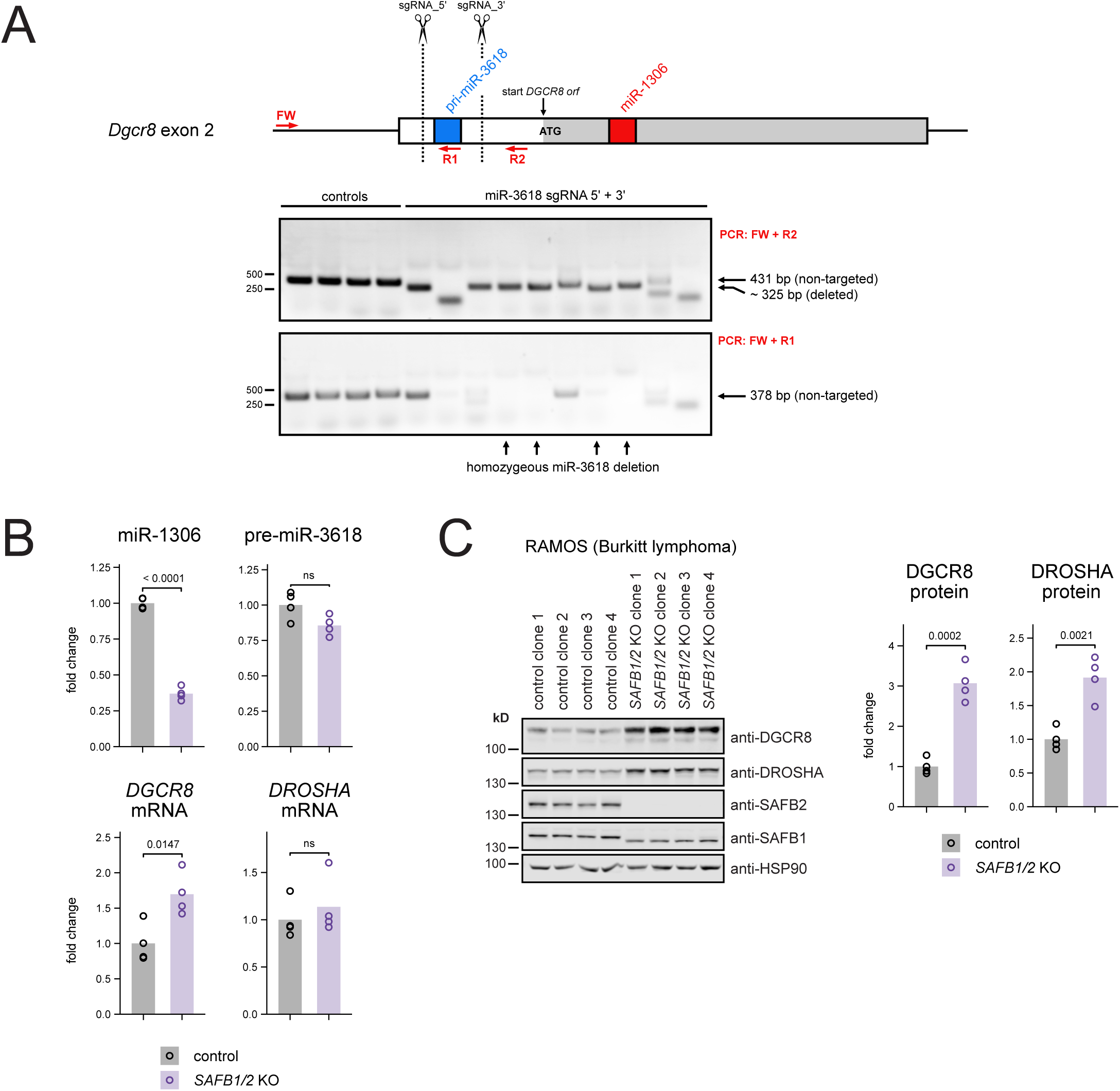
**A**. Schematic drawing of DGCR8 exon 2. Regions encoding pri-miR-3618 and 1306 are shown as blue and red boxes, respectively. SgRNA-targeted sites are marked by dashed lines, primers used for PCR are shown as red arrows. Stained gels show an exemplary PCR analysis of genomic DNA isolated from four controls and 10 clones transiently transfected with Cas9 and flanking sgRNAs (sgRNA_5’ and sgRNA_3’). Primer combinations as indicated. Clones with homozygeous deletion of the miR-3618-encoding sequence are labeled with black arrows. **B**. Quantification of pre-miR-3618, miR-1306, DGCR8 and DROSHA transcript levels in control and SAFB1/2-deficient RAMOS cells by qPCR. Bar charts depict fold changes compared to the control, with each data corresponding to one individual clone. Numbers indicate p-values. **C**. Western blot and bar charts showing protein expression of DGCR8, DROSHA, SAFB1 and SAFB2 for cells as in B. Note that KO clones retain expression of a truncated form of SAFB1, possibly indicating an essential cellular function, but for simplicity are referred to as SAFB1/2-deficient.

**Figure S3.**
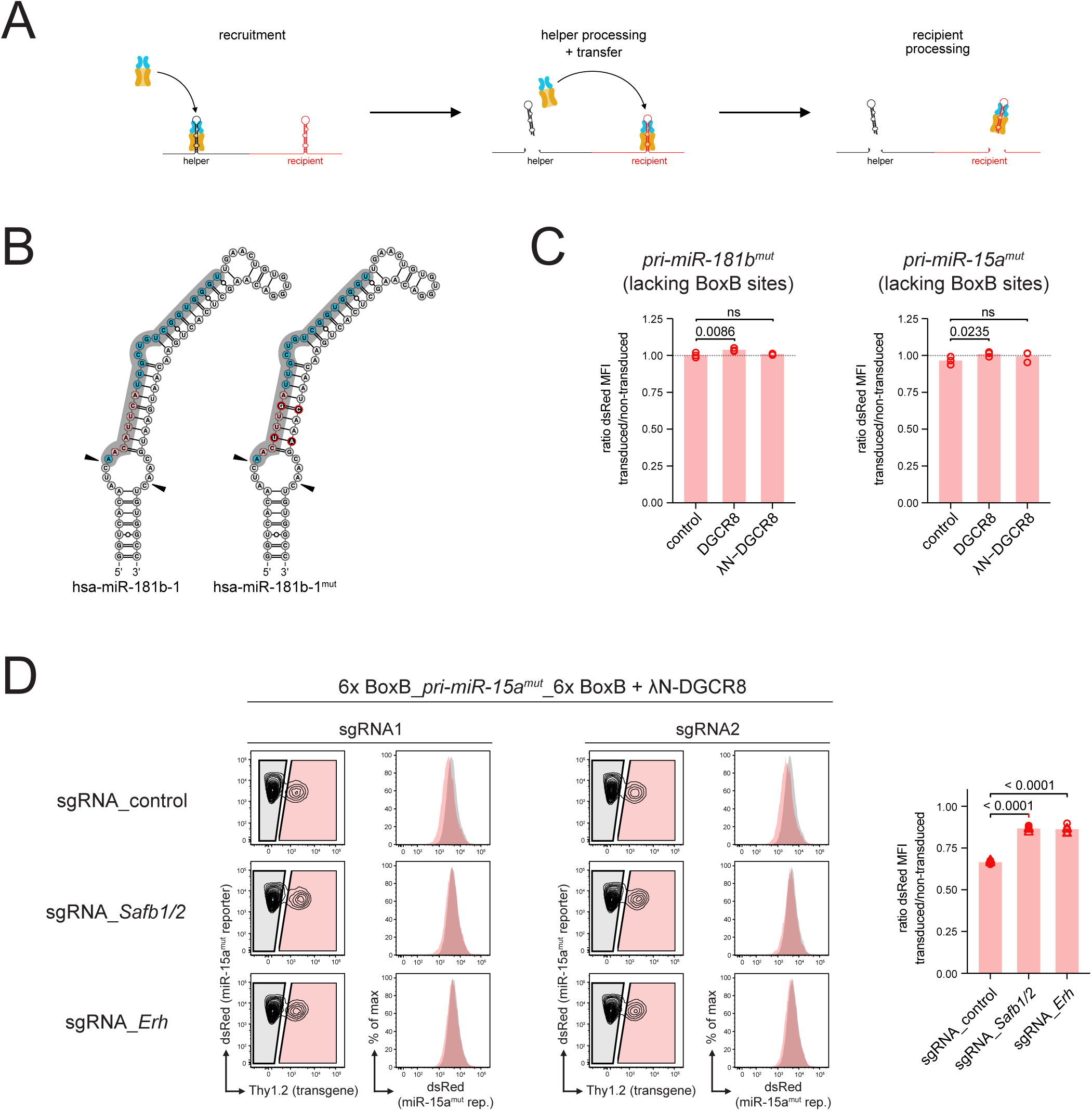
**A**. Model for the temporal sequence of cluster assistance, beginning with Microprocessor recruitment to the helper, followed by helper processing and subsequent transfer of DROSHA/DGCR8 to the recipient. **B**. Predicted structure of the cluster assistance-recipient pri-miR-181b and its seed mutant used for the tethering experiments. The embedded mature miRNA sequences are illustrated as turquise bases over a gray background, with seed sequences in red and black (mutated). Triangles mark the predominant Microprocessor cleavage sites.**C**. Control experiments for the Microprocessor tethering as in Figs. 4B and C, indicating no miRNA reporter repression in the absence on any BoxB aptamer sites in the substrate RNA. **D**. Corresponding experiment to Fig. 4D, using tethering of λN-DGCR8 to miR-15a^mut^ and its respective reporter as a readout. Histogram overlays and the bar graph depict changes in dsRed fluorescence in consequence of SAFB1/2 and ERH knockout. Data points are shown as circles and triangles for the individual sgRNAs (n=3+3). Numbers indicate p-values.

**Figure S4.**
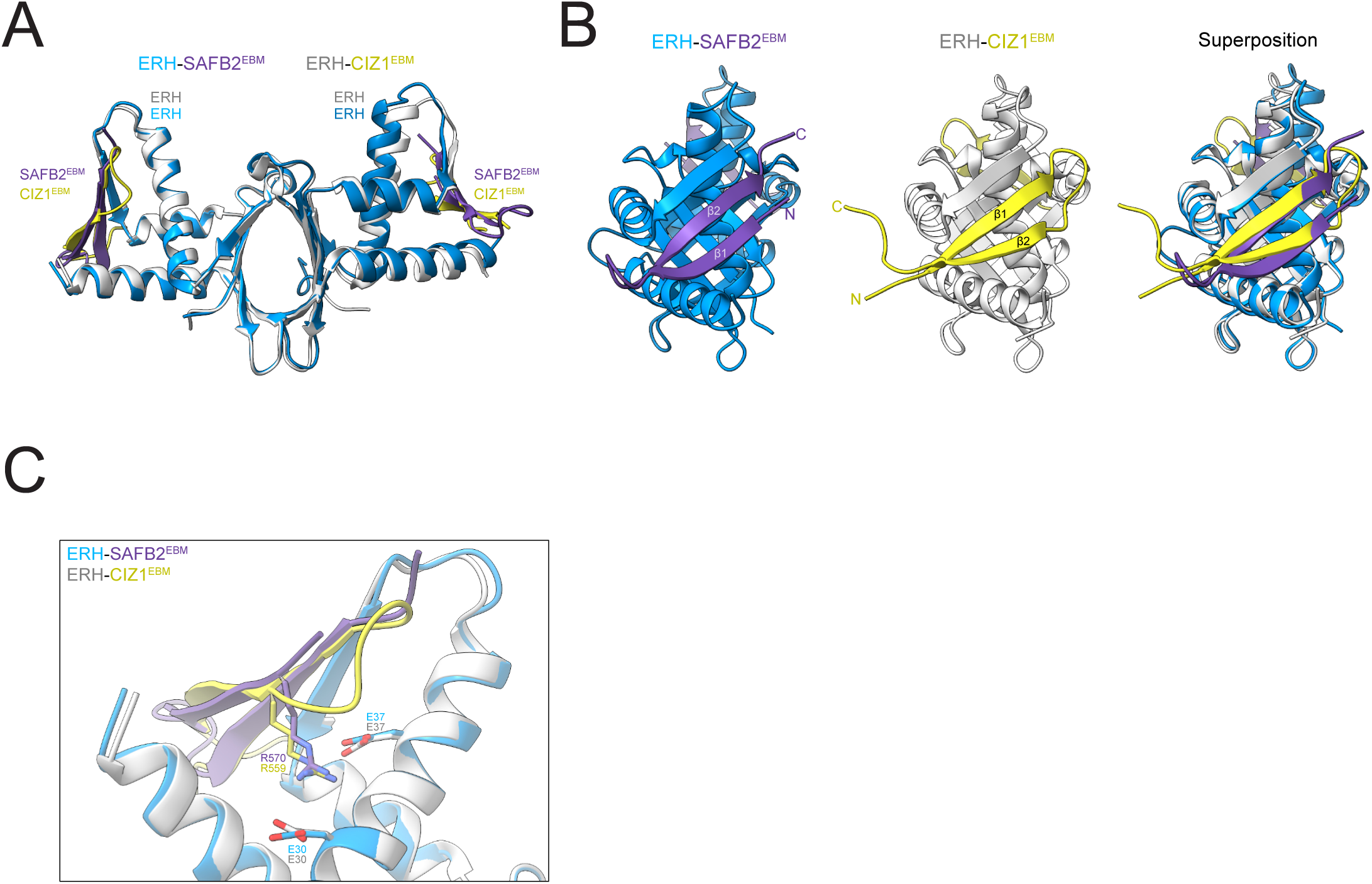
**A, B**. Superposition of the crystal structures of the CIZ1/ERH (pdb: 7×39) and the SAFB2^EBM^/ERH complex, front and side view. **C**. Positioning of critical ERH E30/E37 and SAFB2 R570 as well as CIZ1 R559 residues at the binding interfaces.

**Figure S5.**
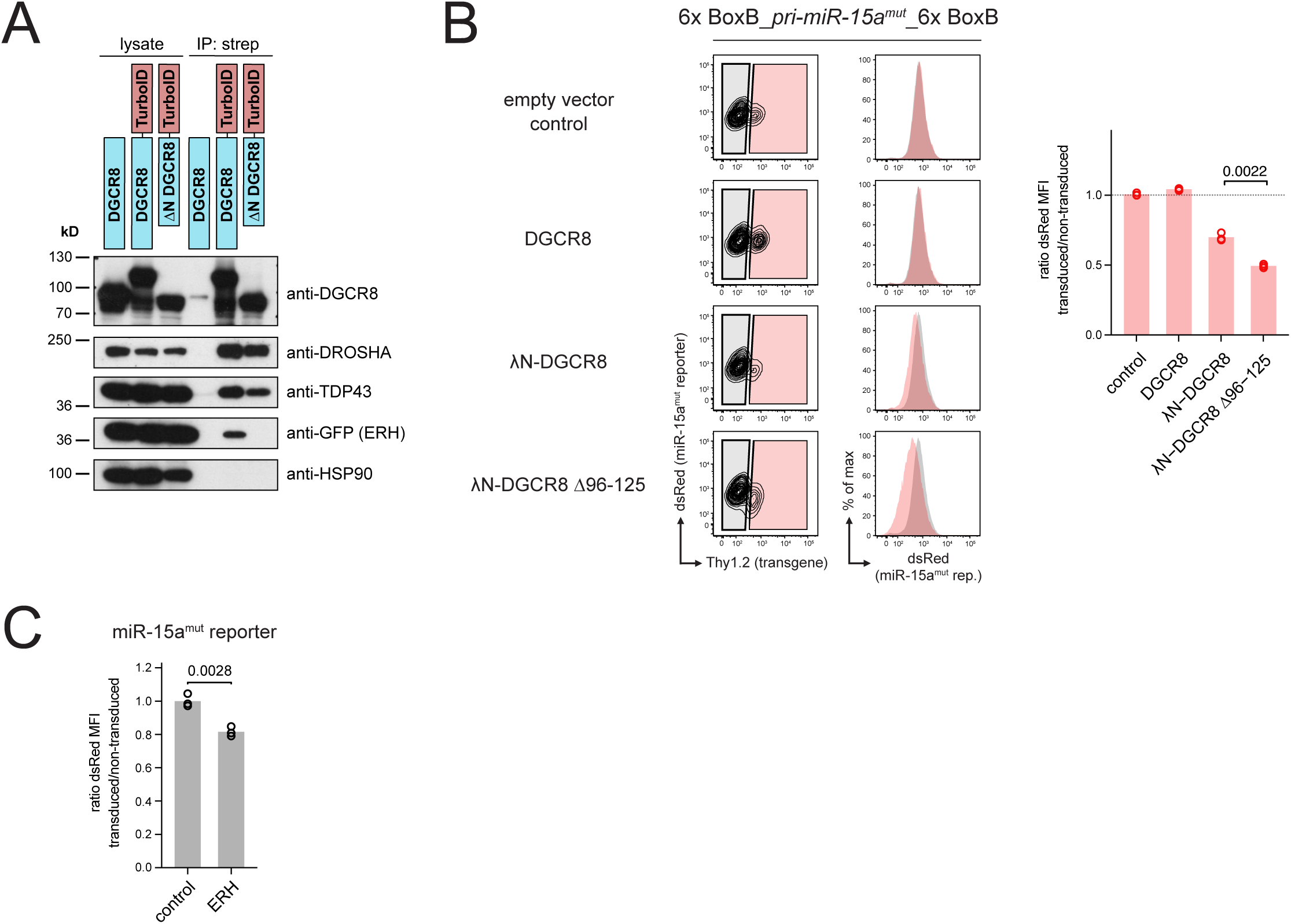
**A**. Western blot analysis of a proximity biotinylation experiment with the indicated TurboID fusion constructs. After labeling, biotinylated proteins were isolated by streptavidin pulldown, separated by SDS-PAGE and blotted. **B, C**. Corresponding experiments to Figs. 5C and D, using miR-15a^mut^ and its respective reporter. Individual bars depict the ratio of miRNA reporter expression in transduced and non-transduced cells, respectively (n=3). Numbers indicate p-values.

**Figure S6.**
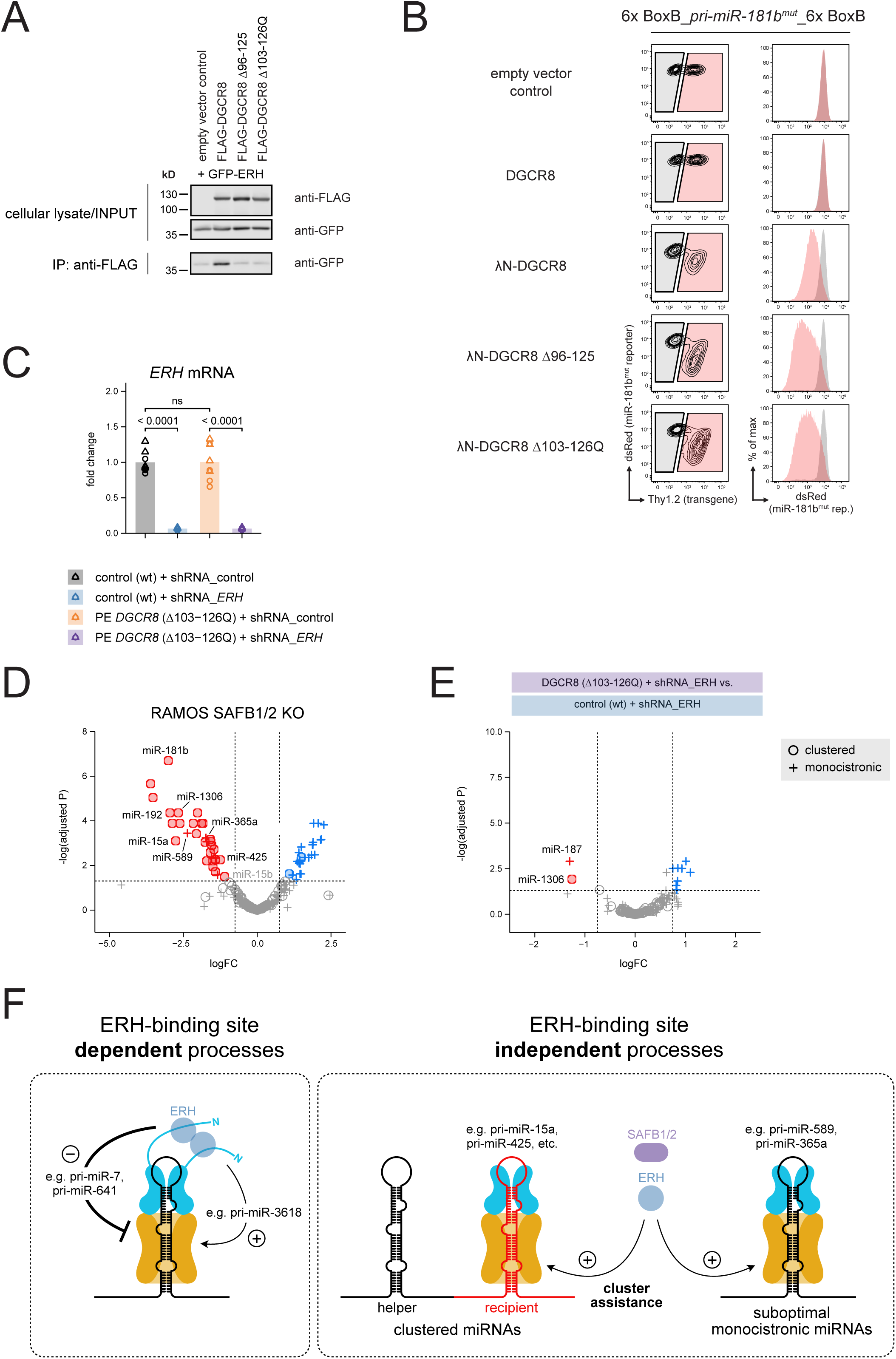
**A**. Western blot analysis of a pulldown experiment in HEK293T cells expressing GFP-ERH and comparing wt DGCR8 with the prime-edited Δ103-126Q variant. Co-immunoprecipitation of the GFP-ERH fusion protein is detected with an anti-GFP antibody. **B**. Flow cytometric analysis of miR-181b^mut^ activity upon tethering of wt DGCR8 and both ERH binding site deletion variants (Δ96-125 and Δ103-126Q) to the BoxB-flanked substrate. Note that both mutants display increased reporter repression compared to the λN-DGCR8 wt control. **C**. Analysis of ERH transcript levels in the individual sample groups as defined in Fig. 6E. Cells were sorted for high expression of the shRNA constructs at d5 post transfection, followed by preparation of total RNA, cDNA synthesis and qPCR. The individual shRNAs (n=4+4) in the respective sample groups are depicted as circles and triangles, respectively. **D**. Volcano plot comparing mature miRNA expression of RAMOS SAFB1/2 KO to control cells (GSE141098; Ref. 31). Vertical dashed lines mark a log-fold change of “-0,75” and “0,75”, respectively, the horizontal line an adjusted p-value <0.05. Down-and upregulated miRNAs are depicted in red or blue, respectively, and either with circles for clustered (defined as with a neighbor on the same strand in less than 2 kb distance) or crosses for monocistronic miRNAs. **E**. Volcano plot as in Fig. 6F, comparing mature miRNA levels between DGCR8 Δ103-126Q and wt clones upon ERH knockdown. **F**. Proposed model for the dual role of ERH in miRNA biogenesis and cluster assistance. Direct ERH binding to DGCR8 has a predominantly inhibitory effect on Microprocessor-mediated cleavage of pri-miRNAs, presumably by bringing the two DGCR8 N-termini together. Beyond that, ERH promotes processing of monocistronic and clustered suboptimal recipients independent of its direct Microprocessor association via the reported binding motif.

## Notes

### Competing Interest Statement

The authors have declared no competing interest.

### Summary of Updates

The manuscript has been re-structured and re-written. The most significant changes include the incorporation of novel small RNA sequencing data for Figs. 1 and 6, and the accompanying updates in the supplementary files.

