## Supplementary Table S3 for "Dual function of ERH in primary miRNA biogenesis"

**Table S2 Data collection and refinement statistics (molecular replacement)**

| | ERH-SAFB2$^{EBM}$ (PDB: 9RMV) |
|---|---|
| **Data collection** | |
| Space group | P 1 21 1 |
| Cell dimensions | |
| *a*, *b*, *c* (Å) | 31.80, 78.48, 53.84 |
| a, b, g (°) | 90.00, 107.05, 90.00 |
| Resolution (Å) | 43.04 - 1.70 (1.77 – 1.70)* |
| No. of reflections | 104,147 (10,990) |
| No. of unique reflections | 31,406 (3,368) |
| $R_{merge}$ (%) | 8.1 (129.9) |
| $R_{pim}$ (%) | 5.3 (84.1) |
| $I$ / s$I$ | 7.7 (0.53) |
| Completeness (%) | 99.9 (99.9) |
| Redundancy | 3.3 (3.3) |
| $CC_{1/2}$ (%) | 99.9 (31.6) |
| **Refinement** | |
| Resolution (Å) | 31.21 – 1.70 (1.77 – 1.70) |
| No. reflections used in refinement | 27,762 (3,062) |
| $R_{work}$ / $R_{free}$ | 18.3 / 20.6 |
| No. atoms | |
| Protein | 2,050 |
| Ligand/ion | 78 |
| Water | 68 |
| *B*-factors | (Ask for input) |
| Protein | 34.43 |
| Ligand/ion | 45.03 |
| Water | 36.38 |
| R.m.s. deviations | |
| Bond lengths (Å) | 0.003 |
| Bond angles (°) | 0.68 |
| Ramachandran favored (%) | 98.74 |
| Ramachandran outliers (%) | 0.00 |

One crystal was used for determining the structure. Highest resolution shell shown in parentheses.

**Table 2** Data collection, phasing and refinement statistics (MIR)

| | Crystal 1 name | Crystal 2 name |
|---|---|---|
| **Data collection** | | |
| Space group | | |
| Cell dimensions | | |
| *a*, *b*, *c* (Å) | | |
| α, β, γ (°) | | |
| Resolution (Å) | ##(high res shell) * | |
| $R_{sym}$ or $R_{merge}$ | ##(high res shell) | |
| $I / \sigma I$ | ##(high res shell) | |
| Completeness (%) | ##(high res shell) | |
| Redundancy | ##(high res shell) | |
| **Refinement** | | |
| Resolution (Å) | | |
| No. reflections | | |
| $R_{work} / R_{free}$ | | |
| No. atoms | | |
| Protein | | |
| Ligand/ion | | |
| Water | | |
| *B*-factors | | |
| Protein | | |
| Ligand/ion | | |
| Water | | |
| R.m.s deviations | | |
| Bond lengths (Å) | | |
| Bond angles (°) | | |

*Number of xtals for each structure should be noted in footnote. *Values in parentheses are for highest-resolution shell.

[AU: Equations defining various *R*-values are standard and hence are no longer defined in the footnotes.]
[AU: Phasing data should be reported in Methods section.]
[AU: Ramachandran statistics should be in Methods section at end of Refinement subsection.]
[AU: Wavelength of data collection, temperature and beamline should all be in Methods section.]

**Table 3** Data collection, phasing and refinement statistics for MAD (SeMet) structures

| | Native | Crystal 1 name | | | Crystal 2 name | |
|---|---|---|---|---|---|---|
| **Data collection** | | | | | | |
| Space group | | common # | | | common # | |
| Cell dimensions | | | | | | |
| *a*, *b*, *c* (Å) | | common # | | | common # | |
| α, β, γ (°) | | common # | | | common # | |
| | *Peak* | *Inflection* | *Remote* | *Peak* | *Inflection* | *Remote* |
| Wavelength | # | # | # | # | # | # |
| Resolution (Å) | # | # | # | # | # | # |
| $R_{sym}$ or $R_{merge}$ | # | # | # | # | # | # |
| $I / \sigma I$ | # | # | # | # | # | # |
| Completeness (%) | # | # | # | # | # | # |
| Redundancy | # | # | # | # | # | # |
| **Refinement** | | | | | | |
| Resolution (Å) | | common # | | | common # | |
| No. reflections | | | | | | |
| $R_{work}$ / $R_{free}$ | | | | | | |
| No. atoms | | | | | | |
| Protein | | | | | | |
| Ligand/ion | | | | | | |
| Water | | | | | | |
| *B*-factors | | | | | | |
| Protein | | | | | | |
| Ligand/ion | | | | | | |
| Water | | | | | | |
| R.m.s deviations | | | | | | |
| Bond lengths (Å) | | | | | | |
| Bond angles (°) | | | | | | |

*Number of xtals for each structure should be noted in footnote. *Values in parentheses are for highest-resolution shell.

[AU: Equations defining various *R*-values are standard and hence are no longer defined in the footnotes.]
[AU: Phasing data should be reported in Methods section.]
[AU: Ramachandran statistics should be in Methods section at end of Refinement subsection.]
[AU: Wavelength of data collection, temperature and beamline should all be in Methods section.]
